# Structural basis of pro-inflammatory signaling via the Interleukin-36 receptor (IL-36R) mediated by IL-36γ and IL-37

**DOI:** 10.1101/2025.02.22.639629

**Authors:** Julie Andries, Martin Toul, Jan Felix, Danielle M. Clancy, Savvas N. Savvides

## Abstract

Interleukin-36 receptor (IL-36R) is activated by IL-36α, IL-36β, and IL-36γ to elicit pro-inflammatory signaling and is targeted in acute skin inflammation by the approved antibody spesolimab. IL-37 was recently proposed as a new IL-36R agonist. Such diverse agonist repertoire together with the antagonistic IL-36Ra and IL-38 create a fascinating structure-function landscape for IL-36R, albeit one that is poorly understood. Here, we elucidate how IL-36R grapples IL-36γ with low affinity to enable facile recruitment of the shared receptor IL-1RAcP with high-affinity. In contrast, IL-36R interacts with IL-37 via the exact opposite binding signature. Comparative interrogation of IL-36R activation by IL-36γ and IL-37 confirmed their common pro-inflammatory signature and distinguished IL-36γ as markedly more pro-inflammatory. Structural comparisons of cytokine-activated versus spesolimab-antagonized IL-36R revealed spesolimab’s mode of action as an allosteric antagonist. Collectively, our study provides the structural and mechanistic blueprint of IL-36R activation by distinct cytokines and will facilitate its therapeutic targeting.

**One Sentence Summary:** Structural blueprint for IL-36R activation by cognate cytokines and its antagonism by spesolimab in generalized pustular psoriasis.

## INTRODUCTION

Interleukin-1 (IL-1) family cytokines are apical mediators released after infection or injury and play key roles in initiating and amplifying inflammatory responses. Interleukin-36 receptor (IL-36R), a key pro-inflammatory receptor in the IL-1 receptor family, plays a pivotal role in inflammatory diseases, particularly skin diseases (*1–3*). IL-36R specifically binds to IL-36 cytokines, which include three agonists—IL-36α, IL-36β, and IL-36γ—and two antagonists, IL-36 receptor antagonist (IL-36Ra) and IL-38. Pro-inflammatory signaling through IL-36R requires the recruitment of IL-1 receptor accessory protein (IL-1RAcP) (*4*). However, binding of IL-36Ra and IL-38 to IL-36R inhibits this recruitment, blocking downstream signal transduction (*5*, *6*). IL-36R is predominantly expressed in skin keratinocytes, which upon activation stimulates NF-κB and MAPK-driven signaling pathways, leading to the production of a variety of pro-inflammatory cytokines and chemokines. Dysregulated IL-36R signaling has been implicated in multiple psoriasis forms (*7*, *8*), including the rare and severe generalized pustular psoriasis (GPP). Mutations in *IL36RN*, the gene encoding IL-36Ra, have been directly linked to GPP pathogenesis (*9*, *10*).

A notable characteristic of the IL-1 cytokine family is the absence of a signal peptide, which typically facilitates conventional ER-Golgi trafficking. Instead, their extracellular release is usually associated with tissue damage, necrosis, and pyroptosis (*11*). In addition, IL-1 cytokines are synthesized as inactive precursors with N-terminal pro-domains requiring proteolytic processing for full activation (*12*). IL-36α, IL-36β, IL-36γ, and IL-36Ra are activated by processing at specific sites—Lys6, Arg5, Ser18, and Val2, respectively—located nine amino acids upstream of a conserved A-X-Asp motif, where A is an aliphatic residue and X is any residue. Incorrect processing by even one amino acid results in a drastic 1,000- to 10,000-fold reduction in activity, akin to the inactive full-length pro-form (*13*). In the case of IL-36γ, the active cytokine is obtained after processing before Ser18 by neutrophil proteases such as cathepsin G or elastase (*14*).

IL-37 is a recent addition to the IL-1 cytokine family (*15–17*). IL-37 is expressed in humans, primates, and other mammals, but not rodents and exists in five isoforms, among which isoform b is the most studied (*18*). IL-37 was initially proposed to be an anti-inflammatory member of the IL-1 cytokine family via its engagement with IL-18R1 and recruitment of the single immunoglobulin interleukin-1-related receptor (SIGIRR) to inhibit signaling (*19*). Akin to related family members, IL-37 also requires proteolytic processing to unleash its bioactivity. (*20*). Previously, IL-37 was shown to be processed by either caspase-1 after Asp20 (*21*, *22*), albeit very inefficiently and at non-physiological concentrations of caspase-1, or before Val46 as determined by Edman degradation (*18*). Recently, IL-37 was identified as a new pro-inflammatory agonist for IL-36R (*20*). Upon proteolytic processing by enzymes released by myeloid cells, such as neutrophil elastase or cathepsin S, IL-37 exhibits pro-inflammatory activity in vitro in HaCaT cells and primary keratinocytes through IL-36R and in vivo, similar to IL-36γ (*20*). However, the structural and mechanistic basis of this interaction had remained enigmatic.

Due to the strong association between IL-36 family cytokines and inflammatory skin diseases such as psoriasis, IL-36R signaling emerged as an attractive therapeutic target. The recent approval by the U.S. Food and Drug Administration (FDA) of the anti-IL-36R monoclonal antibody spesolimab (SPEVIGO^®^, BI 655130) for the treatment of generalized pustular psoriasis, marked a significant advancement in GPP treatment for both adults and pediatric patients aged 12 years and older (*23*, *24*). The crystal structure of the spesolimab Fab fragment with IL-36R (PDB: 6u6u) revealed that the antibody binding epitope is located at the loop connecting D1 and D2 of IL-36R (*25*, *26*). However, the paucity of structural insights into the IL-36R signaling complex, the mechanism by which spesolimab antagonizes IL-36R signaling remains to be fully characterized.

Thus, elucidating the structural basis of the IL-36γ:IL-36R:IL-1RAcP complex will provide the critical insights necessary to consolidate the plethora of immunological research on the pleiotropy of IL-36R signaling over the last two decades and to open new areas of investigation in fundamental and clinical immunology. In this study, we reveal how IL-36R establishes a signaling assembly with the shared receptor IL-1RAcP mediated by pro-inflammatory IL-36γ. By employing a large array of molecular tools and structure-guided mutagenesis to obtain the comparative interaction blueprint of IL-36γ and IL-37 with IL-36R and their relative roles in pro-inflammatory signaling, we make the surprising discovery that while the two cytokines assembles structurally similar complexes, IL-37 displays a much higher affinity towards IL-36R compared to IL-36γ, which intriguingly translates to a markedly weaker pro-inflammatory signaling amplitude. Collectively, our findings will fuel a more targeted interrogation of the emerging interplay between IL-36γ and IL-37 in IL-36R signaling and will impact our understanding of such signaling outputs in physiology, disease, and therapeutic approaches.

## RESULTS

### IL-36γ exhibits low affinity towards its primary receptor IL-36R

To elucidate the mechanism of IL-36γ receptor engagement, we employed isothermal titration calorimetry (ITC) using mature IL-36γ and the extracellular domains (ECD) of IL-36R and IL-1RAcP (**Fig. 1, A and B**). Before ITC titration experiments, protein samples were separately subjected to size-exclusion chromatography coupled to multi-angle laser light scattering (SEC-MALLS) analysis, confirming highly pure, stable and monodisperse samples (**Fig. S1, A to C**). Protein conjugate analysis of IL-36R and IL-1RAcP shows considerable glycosylation on both receptors (**Fig. S1, A and B**). Indeed, there are 9 and 7 predicted N-glycosylation sites in the IL-36R and IL-1RAcP sequences, respectively.

**Fig. 1.**
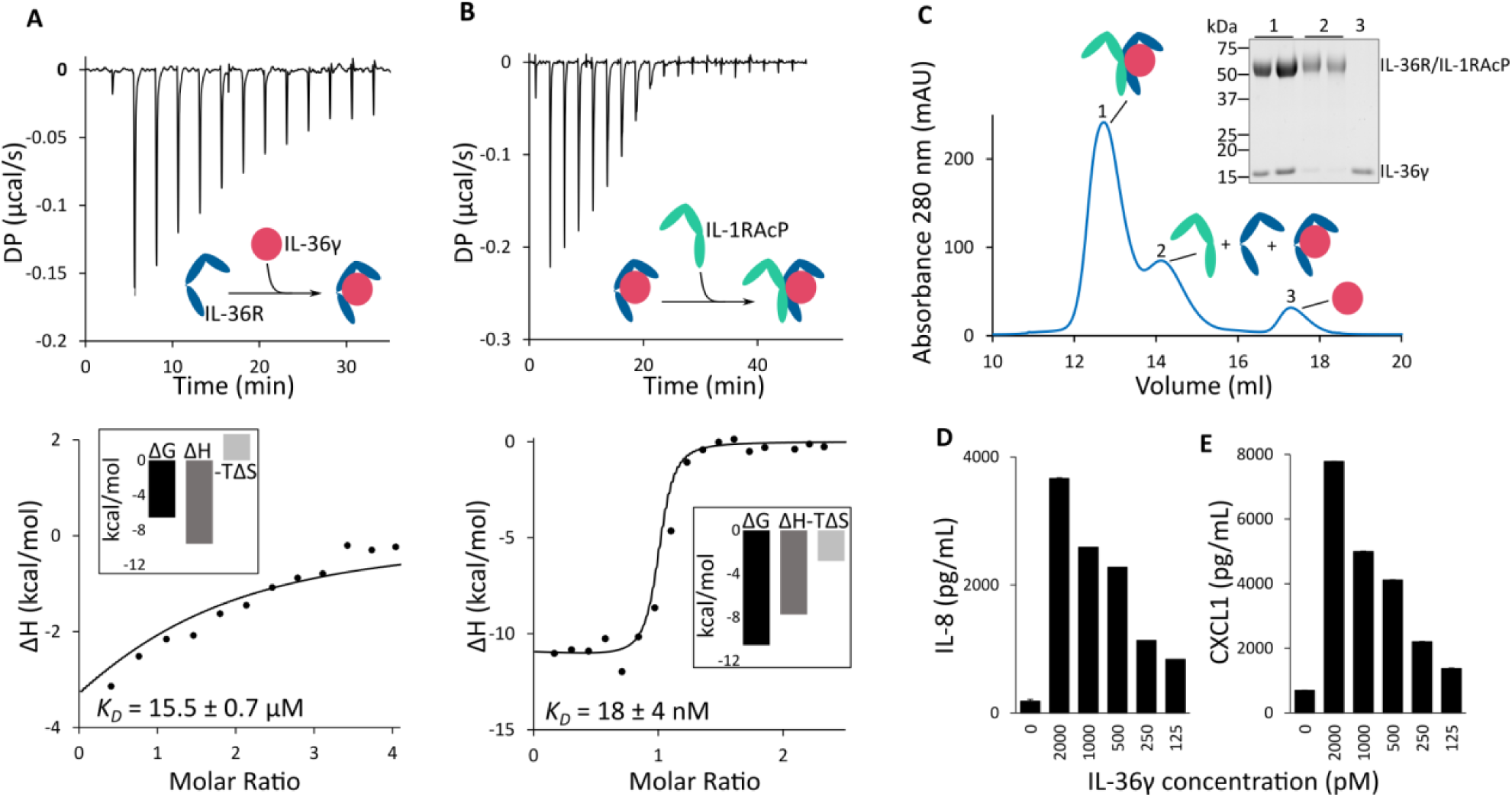
IL-36γ engages IL-36R with low affinity to create a high affinity interface for IL-1RAcP recruitment. (**A-B**) Isothermal titration calorimetry (ITC) experiments of IL-36γ with IL-36R (A), and addition of IL-1RAcP (B) while retaining the sample cell of the binary interaction. A representative thermogram (top) and isotherm (bottom) are displayed for each experiment. A global numerical fitting was performed using KinTek Explorer 10 (KinTek Corporation, USA). Thermodynamic parameters and *K*_D_ ± s.d. values are derived from at least three replicates. Experimental and derived parameters, with corresponding confidence contour analysis, are provided in Table S1 and Fig. S2. Abbreviations: DP, differential electrical power. (**C**) Representative size exclusion chromatogram and SDS-PAGE analysis of IL-36γ ternary complex formation at equimolar concentrations of all components. Peak contents are schematically displayed, where IL-36γ, IL-36R and IL-1RAcP are colored in pink, blue and green, respectively. Sample was injected onto a Superdex 200 Increase 10/300 GL column (Cytiva). (**D-E**) HaCaT cells were incubated overnight at 37°C with mature IL-36γ (Ser18 N-terminus), and IL-8 (D) and CXCL1 (E) concentrations in culture supernatants were determined by ELISA. Error bars represent the percent coefficient of variation (%CV) of triplicate determinations from a representative experiment. Similar results were obtained from three independent experiments.

In contrast to other cytokines in the IL-1 family (*27–30*), ITC experiments showed that IL-36γ exhibits low affinity (*K*_D_ = 15.5 ± 0.7 µM) towards its primary receptor, IL-36R (**Fig. 1A**), consistent with previous surface plasmon resonance (SPR) measurements (*30*). In comparison, IL-1RAcP is recruited with high affinity towards the IL-36γ:IL-36R binary complex (*K*_D_ = 18 ± 4 nM) to assemble the signaling-competent ternary complex (**Fig. 1B**). At the concentrations used, IL-1RAcP showed no apparent binding to either IL-36R or IL-36γ alone, suggesting that a pre-formed complex between IL-36γ and IL-36R is required to create an interface for IL-1RAcP recruitment (**Fig. S1, D to G**).

The low affinity binding of IL-36γ for IL-36R prevented receptor saturation during the binary titration (**Fig. 1A**) and thus likely skewed the binding thermodynamics of ternary complex formation due to ongoing binary complex formation in the latter experimental setup (**Fig. 1B**). Additionally, the apparent binding in ITC titrations did not follow the expected 1:1:1 binding stoichiometry of IL-36γ, IL-36R and IL-1RAcP ternary complex, with low *n*-values (*n* < 1) measured (**Table S1**). Reconstitution of the ternary complex via SEC revealed incomplete complex formation, even at micromolar concentrations of the individual components (**Fig. 1C**), suggesting that a fraction of one of the recombinant complex components is inactive or unavailable for binding. We observed lower *n*-values only in experiments utilizing IL-36R, thus we reasoned this was the source of inactive protein material.

To address these issues, additional analysis of the ITC data was performed using KinTek Explorer 10 (KinTek Corporation, USA) (*31*, *32*). A global numerical fitting was performed with the kinetic model assuming a strict 1:1:1 stoichiometry (*n*=1) and accounting for the low affinity of IL-36γ for IL-36R during the binary titration. This allowed us to uncouple the binding profile of the IL-36γ:IL-36R binary complex interaction from the formation of the IL-36γ:IL-36R:IL-1RAcP complex, allowing more accurate determination of the affinity and thermodynamic parameters of the ternary complex. After uncoupling these two interactions, the affinity of IL-1RAcP for the IL-36γ:IL-36R binary complex was corrected from 44 nM to 18 nM (**Table S1**). The derived parameters, standard errors, and confidence contour analysis using FitSpace Explorer (KinTek Corporation, USA) are presented in Table S1 and Fig. S2 (*33*).

To confirm the biological activity of the recombinant mature IL-36γ used in this study, its activity was tested using the IL-36-responsive keratinocyte HaCaT cell line. In line with previously published results, IL-36γ treated HaCaT cells produced IL-8 and CXCL1 in a dose dependent manner, confirming its pro-inflammatory activity (**Fig. 1, D and E**) (*14*, *20*, *34*).

### Cryo-EM structure of the complete extracellular IL-36R:IL-36γ:IL-1RAcP assembly

To understand how IL-36γ engages its receptor at the molecular level, we determined the structure of the ternary complex comprising IL-36γ bound to the ectodomains of IL-36R and IL-1RAcP by single-particle cryogenic electron microscopy (cryo-EM). The ternary complex was purified via SEC and its composition validated by SDS-PAGE (**Fig. 1C**), mass photometry and Western blot analysis (**Fig. S1, H and I**). To prevent adsorption of the sample on the carbon film and promote optimal particle spreading, the sample was applied to cryo-EM grids in the presence of 0.1 mM dodecyl maltoside (DDM). We collected a total of 8,828 movies, and the resulting 2D classes showed no preferential particle orientations (**Fig. S3**). After iterative rounds of 3D classification, class selection and 3D reconstruction, a final map was obtained with an overall resolution of 3.27 Å according to the gold-standard FSC_0.143_ criterion (**Fig. 2A, Fig. S4, A to C** and **Table S2**).

**Fig. 2.**
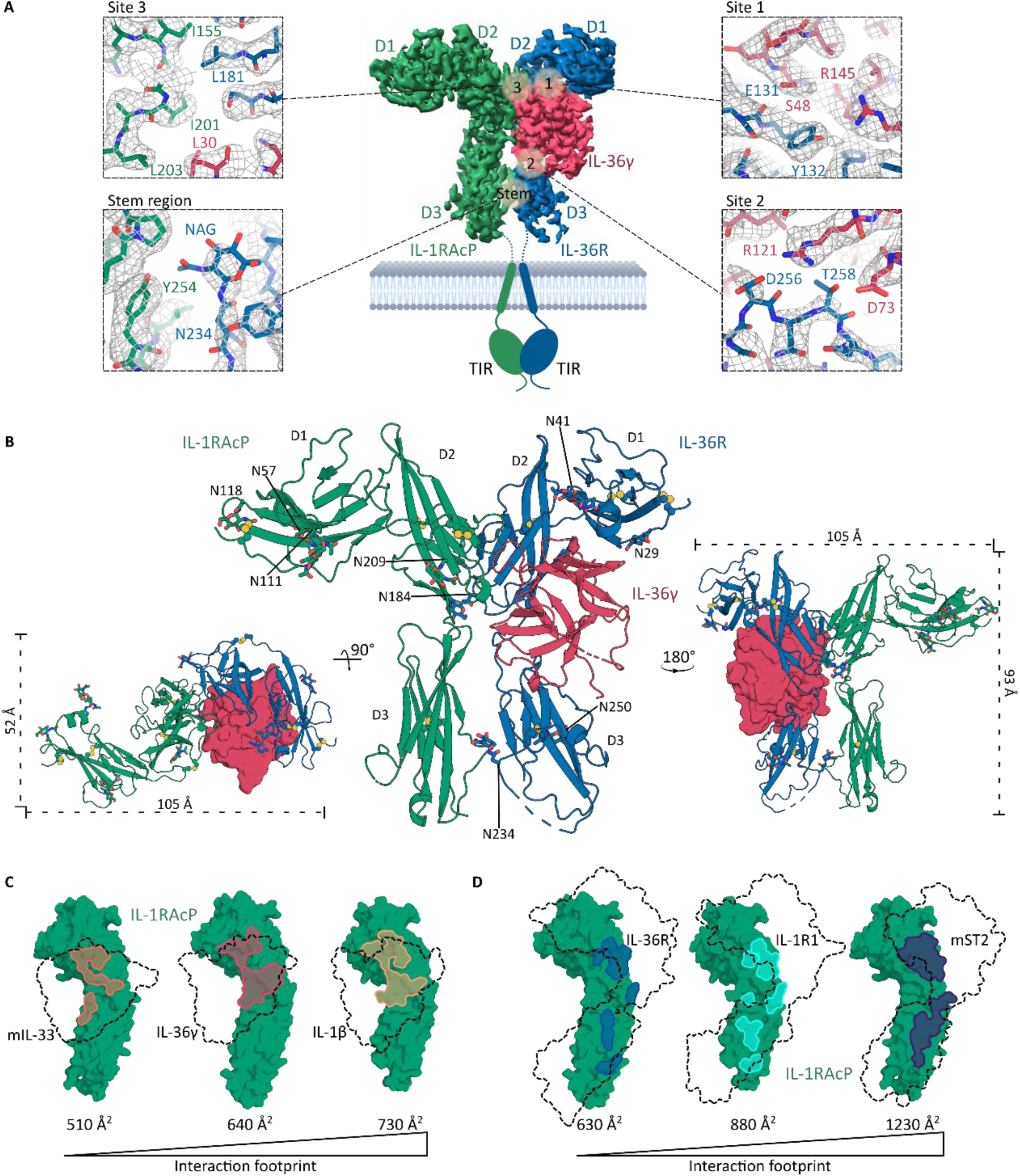
Structural basis for IL-36γ receptor engagement. (**A**) In the middle, a front view of the sharpened 3D map (*B* factor of −70 Å^2^) of the extracellular part of the IL-36γ ternary complex is presented, in combination with a schematic of the transmembrane and TIR domains. The Ig-like domains of the receptors are labeled from D1 to D3. IL-36γ, IL-36R and IL-1RAcP are colored in pink, blue and green, respectively. At the sides, zoom-in views of the different interaction interfaces are shown. The sharpened 3D map is displayed as a grey mesh. (**B**) Top (left), front (middle) and back (right) view of the resulting structural model in cartoon representation, and surface representation for IL-36γ in the top and back view. (**C-D**) Interaction footprints on IL-1RAcP created by IL-1 cytokines (C) or primary receptors (D). Colored patches indicate the interaction interface of human IL-36γ/IL-36R, human IL-1β/IL-1R1 and mouse IL-33/ST2 with IL-1RAcP, based on the cryo-EM structure presented in this study and published crystal structures of the IL-1β complex (PDB: 4dep) and the mouse IL-33 complex (PDB: 5vi4). IL-1RAcP is presented as a green surface, and IL-1 cytokines and primary receptors are shown as a dotted outline. The buried interface area in Å^2^ on IL-1RAcP is noted below the structures. The residues involved in the interfaces are listed in Table S4 and Table S5.

IL-36γ adopts the canonical β-trefoil fold, characteristic of IL-1 family cytokines, and buries approximately 1,400 Å² of surface area upon encapsulation by the three immunoglobulin (Ig) domains of IL-36R (D1, D2, D3) (**Fig. 2B**). The bound form of IL-36γ is nearly identical to the structure of the unbound form, which was previously determined by X-ray crystallography (PDB: 4ize), with a root-mean-square deviation (r.m.s.d.) of 0.501 Å over 122 aligned Cα atoms (*35*). Recruitment of IL-1RAcP buries an extra 1,300 Å² of surface area by interacting with both IL-36γ and IL-36R. The cryo-EM map is well defined across these interaction interfaces, with local resolutions up to 2.8 Å, allowing for confident mapping of interface residues (**Fig. 2A and Fig. S4D**). IL-36γ primarily interacts with D2 and D3 of IL-36R. A flexible linker connects the D1D2 module to the D3 domain of IL-36R (**Fig. 2B**), contributing to the dynamic nature of D3 and resulting in lower local resolution in this area (**Fig. S4D**). In contrast, the relative orientation of D1 and D2 is restricted by a disulfide bridge between Cys118 in the loop connecting D1 and D2 of IL-36R and Cys165 of D2. This disulfide bond is conserved throughout the IL-1 primary receptors, except in IL-1R2, where the loop is replaced by an α-helix (*35–38*).

While the D1 domain of IL-36R is proximal to IL-36γ, the binary interaction is primarily mediated via binding to D2 and D3. These domains engage IL-36γ predominantly through hydrogen bonding, often involving main-chain atoms of either the receptor or cytokine, complemented by the formation of four salt bridges (Glu65, Arg121, Arg126 and Glu164 of IL-36γ with Arg264, Asp256, Asp142 and His137 of IL-36R, respectively) (**Table S3 and Fig. S5**). The prevalence of main-chain interactions may explain IL-36R’s versatility in ligand recognition, despite the lack of sequence conservation among interfacing residues (**Fig. S5**), instead relying on structural homology for ligand binding. This characteristic is also observed in the IL-1β:IL-1R1 complex and to a lesser extent in the IL-33:ST2 complex (*36*, *39*). At the IL-36R D2 interface, Ser48 of IL-36γ interacts with the main chain of IL-36R and Arg145 of IL-36γ forms a hydrogen bond with Tyr132 (**Fig. 2A**, Site 1). At the IL-36R D3 interface, Arg121 of IL-36γ forms a salt bridge with Asp256 and a hydrogen bond with Thr258 of IL-36R (Site 2). The latter also forms a hydrogen bond with Asp73 of IL-36γ. Due to the inherent flexibility of D3 of IL-36R, the interface with IL-36γ is less resolved.

Upon interaction of IL-36γ with IL-36R, IL-1RAcP is recruited to form the signaling-competent ternary complex. Only D2 and D3 of IL-1RAcP are engaged by IL-36γ and IL-36R, while D1 is oriented away from the complex (**Fig. 2B**). At the trimeric junction, a hydrophobic patch is found involving Leu30 and Ala162 of IL-36γ, Leu176 and Leu181 of IL-36R, and Ile155, Ile191, Leu200, Ile201 and Leu203 of IL-1RAcP (**Fig. 2A**, Site 3). In addition to a few hydrogen bonds involving both main-chain and side chains, a single salt bridge is formed between Asp141 of IL-36R and Lys238 of IL-1RAcP. Distal to the cytokine binding interface, heterotypic receptor-receptor interactions involving D3 of both receptors bring the membrane-proximal parts into proximity, facilitating intracellular signaling. One interaction at the stem region involves Tyr254 of IL-1RAcP, forming a hydrogen bond with the N-acetylglucosamine (N-GlcNAc) moiety attached to Asn234 of IL-36R (**Fig. 2A**, Stem). This glycosylation site is conserved in IL-1R1, where the N-GlcNAc moiety on Asn233 of IL-1R1 also forms a hydrogen bond with Tyr254 of IL-1RAcP (*36*). IL-1RAcP is also shared with IL-1R2 and ST2, however neither of these related receptors have an N-glycosylation motif at this position. IL-36R and IL-1RAcP have 9 and 7 Asn-X-Ser/Thr motifs, respectively. In the cryo-EM map, GlcNAc residues were modeled on Asn41, Asn109, Asn184, Asn234 and Asn250 of IL-36R, and Asn57, Asn107, Asn111, Asn118 and Asn209 of IL-1RAcP (**Fig. 2B**).

IL-1RAcP is a shared co-receptor in the IL-1 family, and several crystal structures have been determined for related ternary complexes in this family, including IL-1β:IL-1R1:IL-1RAcP (PDB: 4dep) and mouse IL-33:ST2:IL-1RAcP (PDB: 5vi4) (*36*, *39*). The interface with IL-1RAcP in these structures was extensively compared with our cryo-EM structure using PDBePISA analysis (*40*) and sequence alignments (**Fig. S6**) to compare IL-1RAcP engagement and interaction footprints between these receptor complexes. A comparison of the interface areas of the cytokines on IL-1RAcP reveals that mIL-33 exhibits the smallest interaction interface, followed by IL-36γ and IL-1β (**Fig. 2C**). At the sequence level, IL-1β has more extensive interactions with IL-1RAcP, combining a hydrophobic patch with multiple hydrogen bonds and a salt bridge (**Table S4**). The IL-36γ interface features a similar hydrophobic patch but includes only three hydrogen bonds and no salt bridges. In contrast, the mIL-33 interface has a limited hydrophobic patch, complemented by several hydrogen bonds and a salt bridge. On the receptor-receptor side, the mST2 interaction surface is the most extensive (**Fig. 2D**). This observation is supported by sequence analysis, which reveals numerous hydrogen bonds and salt bridges alongside a hydrophobic patch (**Table S5**). The IL-1R1 interface features several polar and hydrophobic interactions, albeit to a lesser extent, while the IL-36R interface has the most limited set of interactions. Collectively, the IL-36γ:IL-36R binary complex engage IL-1RAcP with fewer interactions compared to these related complexes, which may explain the lower affinity of IL-1RAcP towards IL-36γ:IL-36R (*K*_D_ = 18 ± 4 nM) compared to IL-1β:IL-1R1 (*K*_D_ = 0.9 ± 0.2 nM) or IL-33:ST2 (*K*_D_ = 6.6 ± 3.6 nM) (*39*).

### Spesolimab acts as an allosteric antagonist to inhibit IL-36R signaling

Given the involvement of its activation in several inflammatory skin diseases, IL-36R represents a promising therapeutic target. Patients carrying hypomorphic mutations in IL-36Ra suffer from a rare and severe form of psoriasis termed generalized pustular psoriasis (*41*). Recently, the IL-36R targeting antibody spesolimab (BI 655130) from Boehringer Ingelheim, marketed as SPEVIGO^®^, was approved by the FDA for GPP flares in adults and pediatric patients aged 12 years and older (*23*, *24*). The crystal structure of a Fab fragment of spesolimab in complex with domains D1 and D2 of IL-36R (PDB: 6u6u) revealed that the binding epitope is situated at the loop connecting D1 and D2 (*25*, *26*) (**Fig. 3, A and B**). This binding epitope avoids significant overlap with the IL-36γ or IL-1RAcP interaction sites, making it difficult to rationalize the potent inhibition of IL-36R signaling by spesolimab. As mentioned earlier, the relative orientation of D1 and D2 is restricted due to a conserved disulfide bridge. However, the loop connecting D1-D2 in IL-36R is relatively long compared to other IL-1 family receptors. In IL-36R, this loop consists of 19 amino acids, of which ten are not visible in our cryo-EM map (D119-S129), most likely due to high flexibility. Upon binding of the spesolimab Fab fragment, this loop is pushed inwards toward the cytokine-binding interface, resulting in a clash with the β3-β4 loop of IL-36γ, according to our cryo-EM structure of the IL-36γ:IL-36R:IL-1RAcP complex (**Fig. 3C**) (*26*). This suggests that the allosteric binding of spesolimab would be competitive with respect to cytokine engagement.

**Fig. 3.**
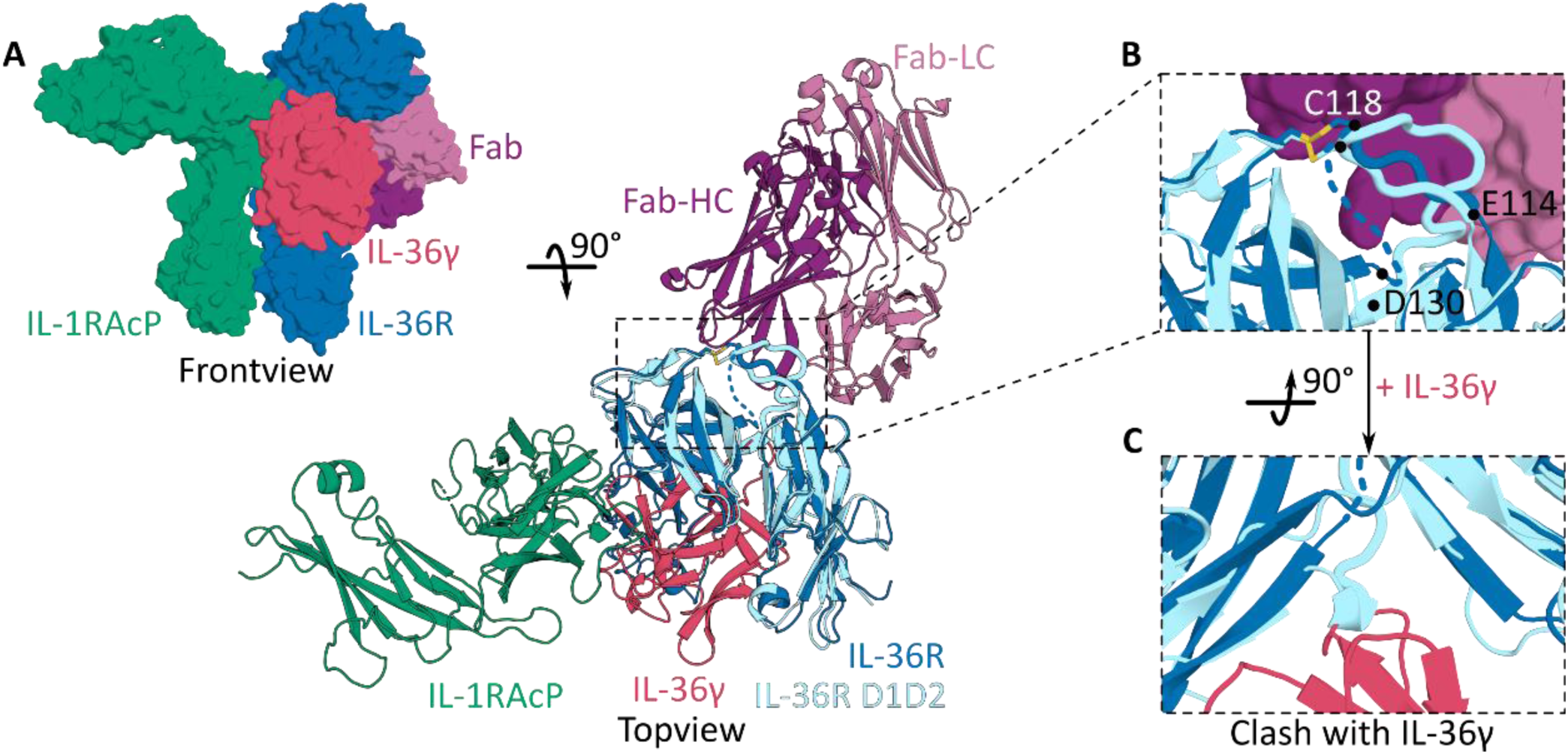
Structural basis for the allosteric competitive binding mode of spesolimab. (**A**) Front view and top view of the cryo-EM structure of the IL-36γ ternary complex aligned with the crystal structure (PDB: 6u6u) of Fab BI 655130 (spesolimab) with IL-36R domain 1 and 2. (**B**) Zoom-in view of the Fab fragment interacting with the loop connecting IL-36R D1 and D2 (IL-36R in light blue), aligned with IL-36R from our cryo-EM structure (IL-36R in dark blue), where the loop is partially unresolved in the density (dotted line). (**C**) Zoom-in at the IL-36γ interface, showing a clash with the protruding D1-D2 loop after binding of the Fab. Abbreviations: HC, heavy chain; LC, light chain.

### IL-37 exhibits a higher affinity for IL-36R than IL-36γ

IL-37 was recently identified as a novel pro-inflammatory agonist of IL-36R (*20*), however the biophysical characteristics of this interaction have not been fully elucidated. As previously mentioned, IL-1 family cytokines require proteolytic processing to unleash their inflammatory activity. We tested different mature constructs based on literature starting at Val46, His47 and Lys53 (*18*, *20*), and found that only the mature cytokine starting at Lys53 induced robust pro-inflammatory cytokine production in HaCaT cells (**Fig. S7A**). A multiple sequence alignment of IL-37 and the IL-36 cytokines also revealed that Lys53 of IL-37 is located 9 amino acids upstream of the conserved *A-X-Asp* motif (**Fig. S5**) (*13*). This is in agreement with a previous study reporting processing by cathepsin S at this residue (*20*). Therefore, we proceeded with Lys53-IL-37 for further characterization, referring to it as IL-37.

We examined the interaction of IL-37 with the ectodomains of IL-36R using ITC. At the concentrations used for these experiments, IL-37 behaves as a dimer in contrast to IL-36γ and IL-36Ra which behave as monomers, as confirmed by SEC-MALLS (**Fig. S7B**). Previously reported dissociation constants of IL-37 dimerization range between 5 to 294 nM (*21*, *42*). Interestingly, IL-37 exhibits >200-fold higher affinity towards IL-36R than IL-36γ, with a dissociation constant of 68 ± 34 nM for IL-37 compared to 15.5 ± 0.7 µM for IL-36γ (**Fig. 1A** and **Fig. 4A**). Unlike IL-36γ, the interaction of IL-37 with IL-36R is endothermic and mainly entropically driven. As before, we utilized the KinTek Explorer software to account for the inactive fraction of recombinant IL-36R. Surprisingly, upon addition of IL-1RAcP to the IL-37:IL-36R binary complex, no heat of interaction was detected (**Fig. 4B**), even when the concentration of IL-1RAcP was increased up to 427 µM (**Fig. S7C**). As a control, we demonstrated that the titration of IL-1RAcP in IL-37, and IL-37 in buffer, do not produce heat changes (**Fig. S7, D and E**). Nonetheless, IL-37 induced the production of pro-inflammatory cytokines IL-8 and CXCL1 in HaCaT cells, albeit only at concentrations several orders of magnitude higher than required for IL-36γ (**Fig. 4, C and D**, and **Fig. 1, D and E**). For signaling to occur, two receptor chains with functional intracellular TIR domains must be brought into proximity. Previous knockdown experiments have proposed IL-1RAcP as the co-receptor for IL-37 (*20*). Silencing of IL-36R, IL-1RAcP or the downstream signaling adaptor MyD88 by RNA interference suppressed IL-37-induced cytokine production, whereas silencing IL-18R1 or SIGIRR did not (*20*). Given that IL-37 signaling appears to rely on IL-1RAcP, the reduced inflammatory response compared to IL-36γ might be due to lower affinity between IL-1RAcP and the IL-37:IL-36R binary complex, as seen in ITC measurements. Since this interaction was undetectable even at 427 µM (Fig. S7C), the affinity is likely well into the mM range. A similar characterized low-affinity interaction in cytokine biology is the binding of IL-10R2 to the IL-10:IL-10R1 complex, which has a dissociation constant of 234 ± 61 µM (*43*).

**Fig. 4.**
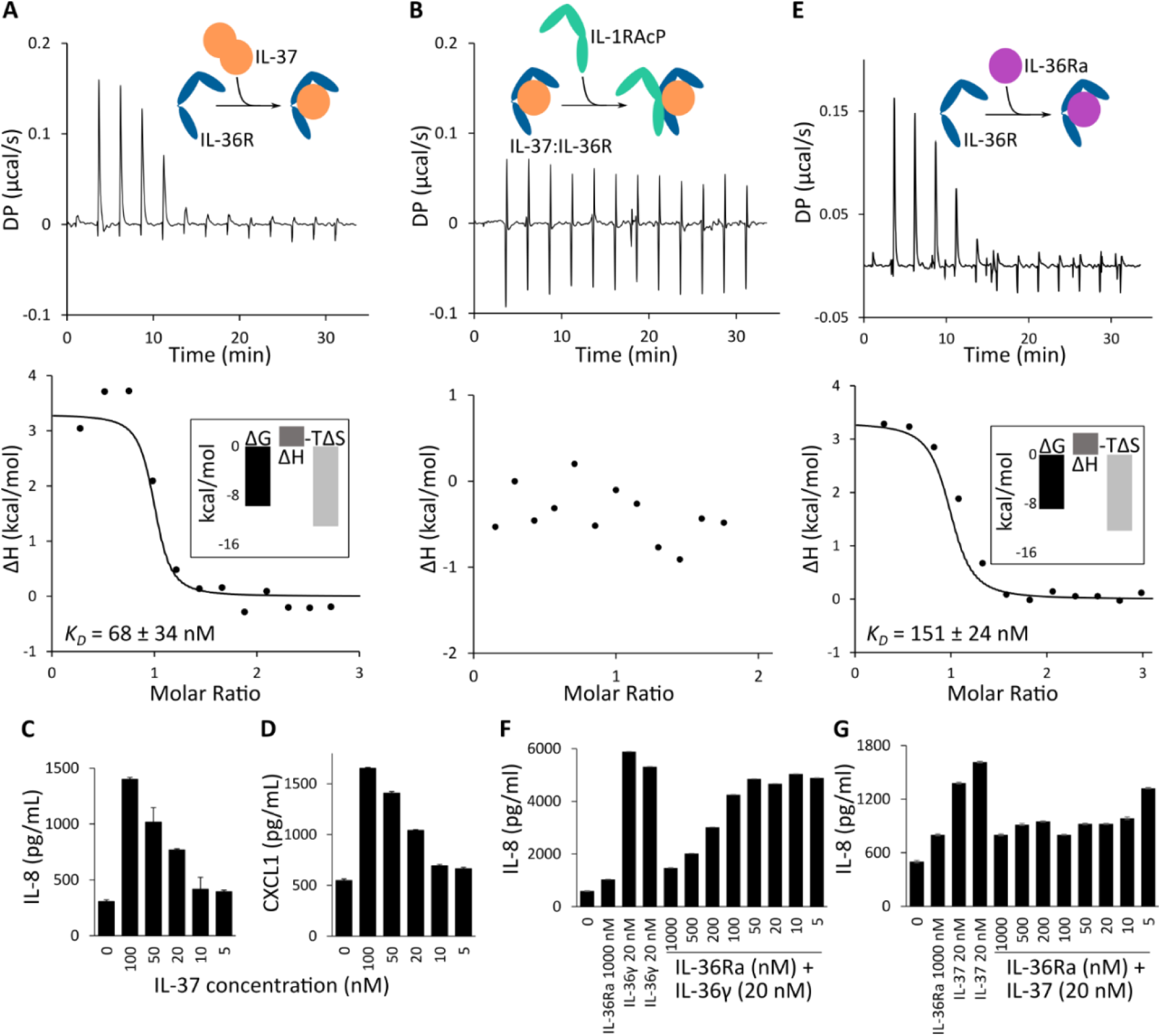
IL-37 binds IL-36R with higher affinity but induces a lower pro-inflammatory output than IL-36γ. (**A-B**) ITC experiments of IL-37 with IL-36R (A), and addition of IL-1RAcP (B) while retaining the sample cell of the binary interaction. A global numerical fitting was performed using KinTek Explorer 10 (KinTek Corporation, USA) and single representative thermograms (top) and isotherms (bottom) are displayed. Thermodynamic parameters and *K*_D_ ± s.d. values are derived from at least three replicates. Experimental and derived parameters, with corresponding confidence contour analysis, are provided in Table S1 and Fig. S2. Abbreviations: DP, differential electrical power. (**C-D**) HaCaT cells were incubated overnight at 37°C with mature IL-37 (Lys53 N-terminus), and IL-8 (C) and CXCL1 (D) concentrations in culture supernatants were determined by ELISA. (**E**) ITC experiment of IL-36Ra with IL-36R. The same analysis procedure, as described for (A-B), was applied. (**F-G**) HaCaT cells were pretreated with IL-36Ra for 1 hour at 37°C at the indicated concentrations, after which 20 nM IL-36γ (F) or IL-37 (G) was added. The day after, the concentration of IL-8 in the cell culture supernatants was measured by ELISA. Error bars represent the percent coefficient of variation (%CV) of triplicate determinations from a representative experiment. Similar results were obtained from three independent experiments.

IL-36Ra is a naturally occurring antagonist of IL-36R, previously reported to interact with IL-36R with higher affinity than the various IL-36 agonists (*30*). Upon confirming the monodispersity and monomeric nature of IL-36Ra by SEC-MALLS and benchmarking its heat of dilution in ITC (**Fig. S7, B and F**), we proceeded to measure a nanomolar affinity of IL-36Ra towards IL-36R with *K*_D_ = 151 ± 24 nM by ITC (Fig. 4E), although this is considerably weaker than the previously reported *K*_D_ = 5.8 ± 0.3 nM measured by SPR (*30*). The interaction between IL-36Ra and IL-36R is endothermic with a strong entropic component dominating the thermodynamic profile, like IL-37 (**Fig. 4E**). As expected, the addition of IL-36Ra suppressed both IL-36γ- and IL-37-induced pro-inflammatory cytokine production in HaCaT cells (**Fig. 4, F and G**).

### Mutational analysis reveals critical IL-37 and IL-36γ interaction sites on IL-36R

IL-37 shares significant structural similarity with IL-36γ, as indicated by r.m.s.d. values of 0.77/0.59 Å over 98 or 89 aligned Cα atoms, respectively, when aligning previously solved crystal structures of IL-37 (PDB codes: 6ncu and 5hn1) with the bound form of IL-36γ from our cryo-EM analysis (*42*, *44*). However, despite their common receptor, the key interfacing residues between IL-37 and IL-36γ are largely non-conserved (**Fig. S5**). Due to the low affinity of the IL-37:IL-36R:IL-1RAcP, we were unable to pursue structural studies of the IL-37 ternary complex by cryo-EM. However, AlphaFold3 generated a high-confidence structural model of the IL-37:IL-36R complex, with an interface predicted TM (ipTM) score exceeding 0.8, suggesting a reliable prediction of the interaction interface (**Fig. S8, A and B**). The model indicates a binding mode for IL-37 to IL-36R similar to that of IL-36γ, as supported by the structural alignment with the extracted IL-36γ:IL-36R complex from our cryo-EM structure, which shows an r.m.s.d. value of 2.03 Å over 386 aligned Cα atoms (**Fig. S8A**). Using structural insights obtained from our cryo-EM structure of the IL-36γ ternary complex, in combination with predicted models, we selected two specific interaction sites to further interrogate the differential engagement and affinity of these distinct cytokines to IL-36R.

At site 1 in the IL-36γ:IL-36R interaction, Ser48 and Ser50 occupy the positions corresponding to Lys83 and Tyr85 in IL-37, with only Ser48 forming a hydrogen bond with the main chain of Glu131 in IL-36R (**Fig. 5A**). In the Alphafold3-predicted complex, the hydrophobic portion of Lys83 in IL-37 is nestled between the aromatic rings of Tyr85 and Tyr132 in IL-36R, while the ε-amino group forms hydrogen bonds with both tyrosine residues. These hydrophobic interactions could partially explain the higher affinity of IL-37 for IL-36R compared to IL-36γ. To further explore this, we mutated Ser48 and Ser50 in IL-36γ to the corresponding residues in IL-37 (IL-36γ_S48K/S50Y_), resulting in an approximately 10-fold increase in affinity for IL-36R compared to wild-type IL-36γ (**Fig. 5B**). Interestingly, a more prominent entropic contribution, similar to that observed in the interaction of IL-37 with IL-36R, was observed. When IL-1RAP was titrated into this binary complex, the affinity was similar to the wild-type complex (**Fig. 5C**).

**Fig. 5.**
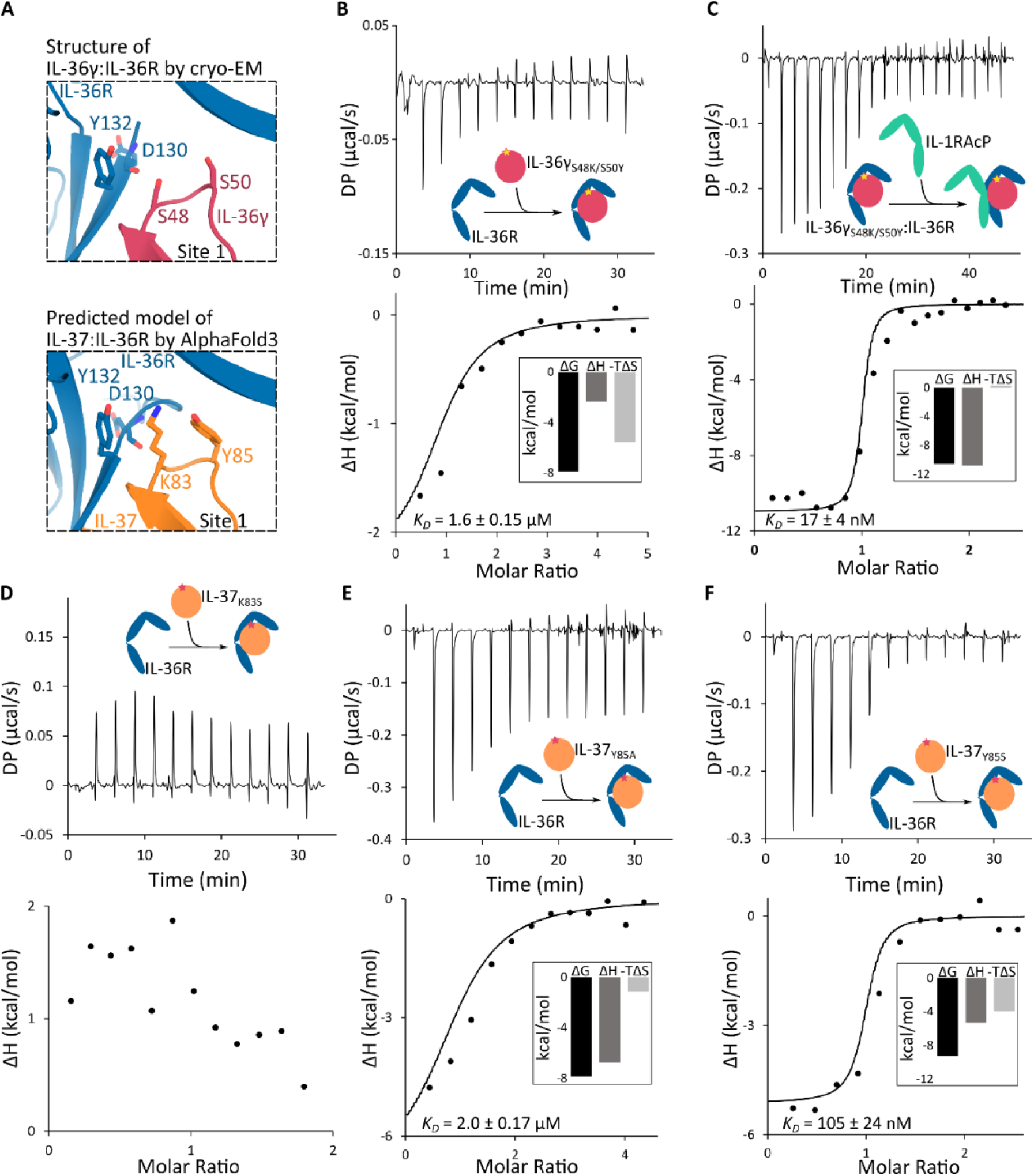
Exchange of residues between IL-36γ and IL-37 at the IL-36R D2 interface. (**A**) Zoom-in views at the IL-36R D2 interface with IL-36γ of the experimental model from our cryo-EM structure (top) and predicted model by AlphaFold3 of IL-37 with IL-36R (bottom) (Fig. S8, A and B). IL-36γ, IL-37 and IL-36R are colored in pink, orange and blue, respectively. (**B-F**) ITC experiments of IL-36γ_S48K/S50Y_ (B), IL-37_K83S_ (D), IL-37_Y85A_ (E) and IL-37_Y85S_ with IL-36R (F), and addition of IL-1RAcP (C) while retaining the sample cell of the binary interaction (B). A global numerical fitting was performed using KinTek Explorer 10 (KinTek Corporation, USA) and single representative thermograms (top) and isotherms (bottom) are displayed. Thermodynamic parameters and *K*_D_ ± s.d. values are derived from at least three replicates. Experimental and derived parameters, with corresponding confidence contour analysis, are provided in Table S1 and Fig. S2. Abbreviations: DP, differential electrical power.

Interestingly, Lys83 and Tyr85 in IL-37 at the site 1 interface have also been identified as crucial residues driving IL-37 dimerization (*42*). At the cytokine dimer interface, Tyr85 of IL-37 is buried within a hydrophobic pocket of the opposing monomer, stabilizing the dimerization. Additionally, Asp73 and Lys83 of form salt bridges at the solvent-exposed edges of this hydrophobic pocket. Both mutation of Tyr85 to alanine (Y85A) and a charge reversal of Asp73 (D73K) completely abolished dimerization (*42*). In line with this, mutating Lys83 and Tyr85 to serine abolished dimerization and resulted in monomeric IL-37 mutant variants, similar to IL-37_Y85A_ (**Fig. S8, D and E**). Surprisingly, the mutation of Lys83 in IL-37 to serine (K83S) abolished detectable interaction with IL-36R (**Fig. 5D**). While the IL-37_Y85S_ variant exhibited a slight decrease in affinity compared to wild-type IL-37, the Y85A mutation led to a more pronounced 34-fold reduction (**Fig. 5, E and F**). Interestingly, both IL-37_Y85S_ and IL-37_Y85A_ variants flipped the isotherm from an endothermic to an exothermic profile (**Fig. 5, E and F**), with a notably reduced entropic contribution, particularly for IL-37_Y85S_. This shift could be attributed to the loss of hydrophobic packing between Tyr85, Lys83, and Tyr132. Additionally, we selected residues for mutagenesis in site 2 at the interface of the cytokine with the D3 domain of IL-36R. In IL-36γ, Arg121 forms a salt bridge and a hydrogen bond with Asp256 and Thr258 in IL-36R, respectively (**Fig. 6A**). This arginine is conserved across IL-36Ra and IL-37, corresponding to Arg102 in IL-36Ra and Arg158 in IL-37. In IL-36Ra, the mutation of this arginine to tryptophan (p.Arg102Trp) has been associated with GPP (*10*, *45*). In this context, the bulky and hydrophobic tryptophan side chain likely disrupts the interaction of IL-36Ra with D3 of IL-36R, impairing its regulatory function. When Arg121 in IL-36γ was substituted with alanine, the loss of the salt bridge and hydrogen bond did not considerably alter its affinity for IL-36R, nor did it affect the affinity of IL-1RAcP for the IL-36γ:IL-36R complex (**Fig. 6, B and C**). In contrast, mutating Arg158 in IL-37 to alanine abrogated any measurable interaction with IL-36R (**Fig. 6, D and E**).

**Fig. 6.**
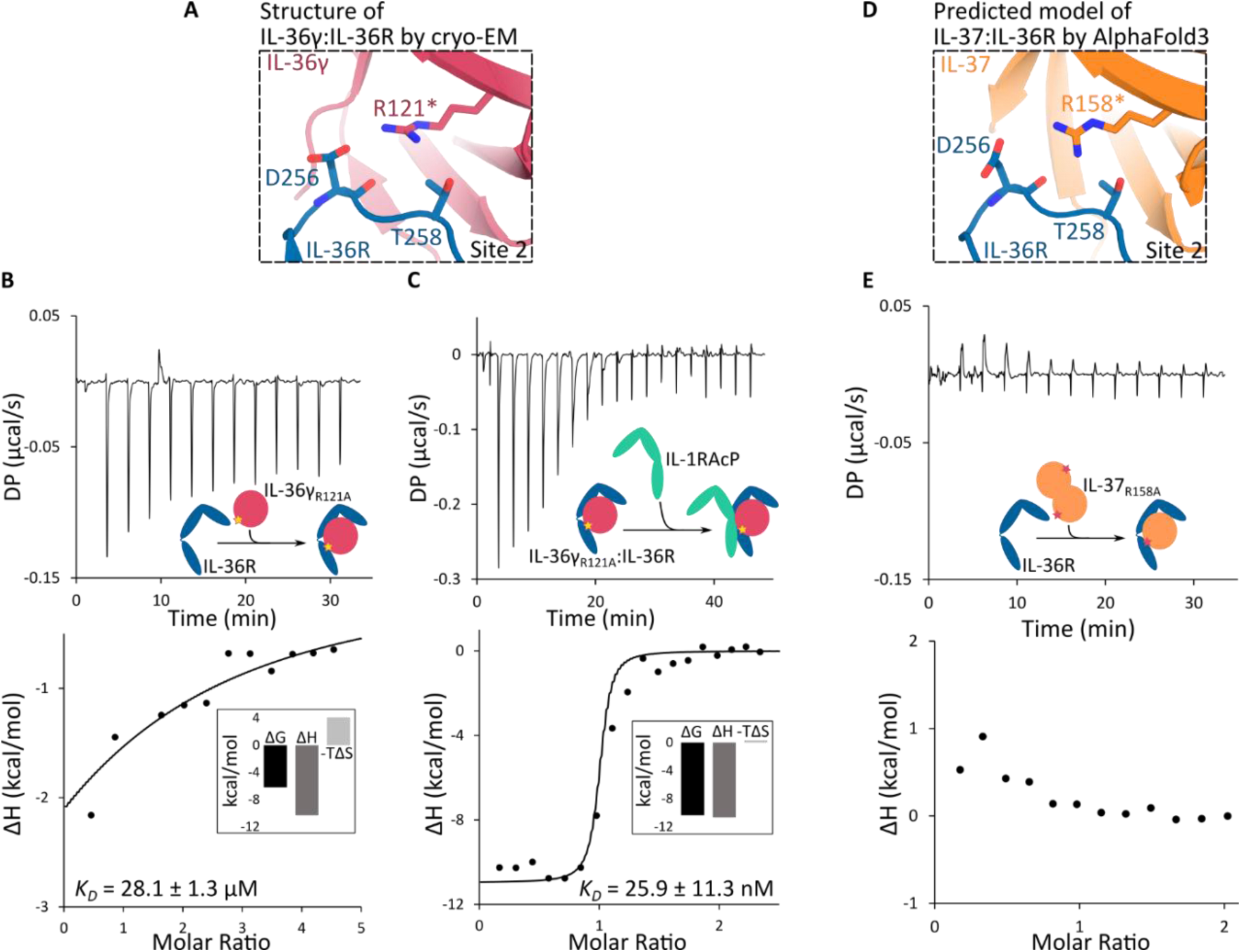
Mutagenesis of the conserved arginine in the cytokines at the interface with IL-36R D3. (**A**) Zoom-in view at the IL-36R D3 interface of the experimental model from our cryo-EM structure. IL-36γ and IL-36R are colored in pink and blue respectively. (**B-C**) ITC experiments of IL-36γ_R121A_ with IL-36R (B), and addition of IL-1RAcP (C) while retaining the sample cell of the binary interaction (B). (**D**) Zoom-in view at the IL-36R D3 interface of the predicted model by AlphaFold3 of IL-37 with IL-36R (Fig. S8, A and B). IL-37 and IL-36R are colored in orange and blue, respectively. (**E**) ITC experiment of IL-37_R158A_ with IL-36R. A global numerical fitting was performed using KinTek Explorer 10 (KinTek Corporation, USA) and single representative thermograms (top) and isotherms (bottom) are displayed. The blue isotherm, as plotted by KinTek Explorer 10, represents the integrated counterpart of the black conventional isotherm. Thermodynamic parameters and *K*_D_ ± s.d. values are derived from at least three replicates. Experimental and derived parameters, with corresponding confidence contour analysis, are provided in Table S1 and Fig. S2. Abbreviations: DP, differential electrical power.

To further investigate the implications of these mutations, we compared the chromatographic behavior and molar mass of the mutants to the wild-type proteins using SEC-MALLS (**Fig. S8, C to E**). IL-36γ typically exists as a monomer, except at very high concentrations, which was also observed in the two IL-36γ mutants, although the small dimer peak became more pronounced in the IL-36γ R121A mutant. The double mutant IL-36γ_S48K/S50Y_ exhibited a slight alteration in chromatographic mobility, yet its molar mass confirmed it as a monomer. As previously mentioned, all three mutations at the dimer interface of IL-37 (K83S, Y85A, Y85S) completely abolished dimerization, consistent with previous findings for Y85A (*42*). Mutating Arg158 to alanine in IL-37, a residue situated away from the dimer interface, did not disrupt dimer formation (**Fig. S8, C to E**).

To assess the impact of these mutations on the pro-inflammatory output, we stimulated HaCaT cells with the different mutant variants and quantified IL-8 and CXCL1 production by ELISA, as before (**Fig. 7**). Although both IL-36γ_S48K/S50Y_ and IL-36γ_R121A_ exhibited similar affinities to IL-36R as wild-type IL-36γ in ITC titrations, only IL-36γ_S48K/S50Y_ maintained comparable pro-inflammatory activity, while IL-36γ_R121A_ displayed reduced activity (**Fig. 7, A and B**). Consistent with ITC results, both IL-37_K83S_ and IL-37_R158A_ did not induce pro-inflammatory output above background levels (**Fig. 7, C and D**). IL-37_Y85A_, which showed much lower affinity for IL-36R in ITC, failed to elicit IL-8 or CXCL1, while IL-37_Y85S_, despite comparable affinity to wild-type, resulted in greatly reduced IL-8 and CXCL1 induction (**Fig. 7, C and D**). In general, the discrepancies between the ITC results and the cellular assays suggest that even a slight decrease in affinity can have a huge effect on the pro-inflammatory outcome.

**Fig. 7.**
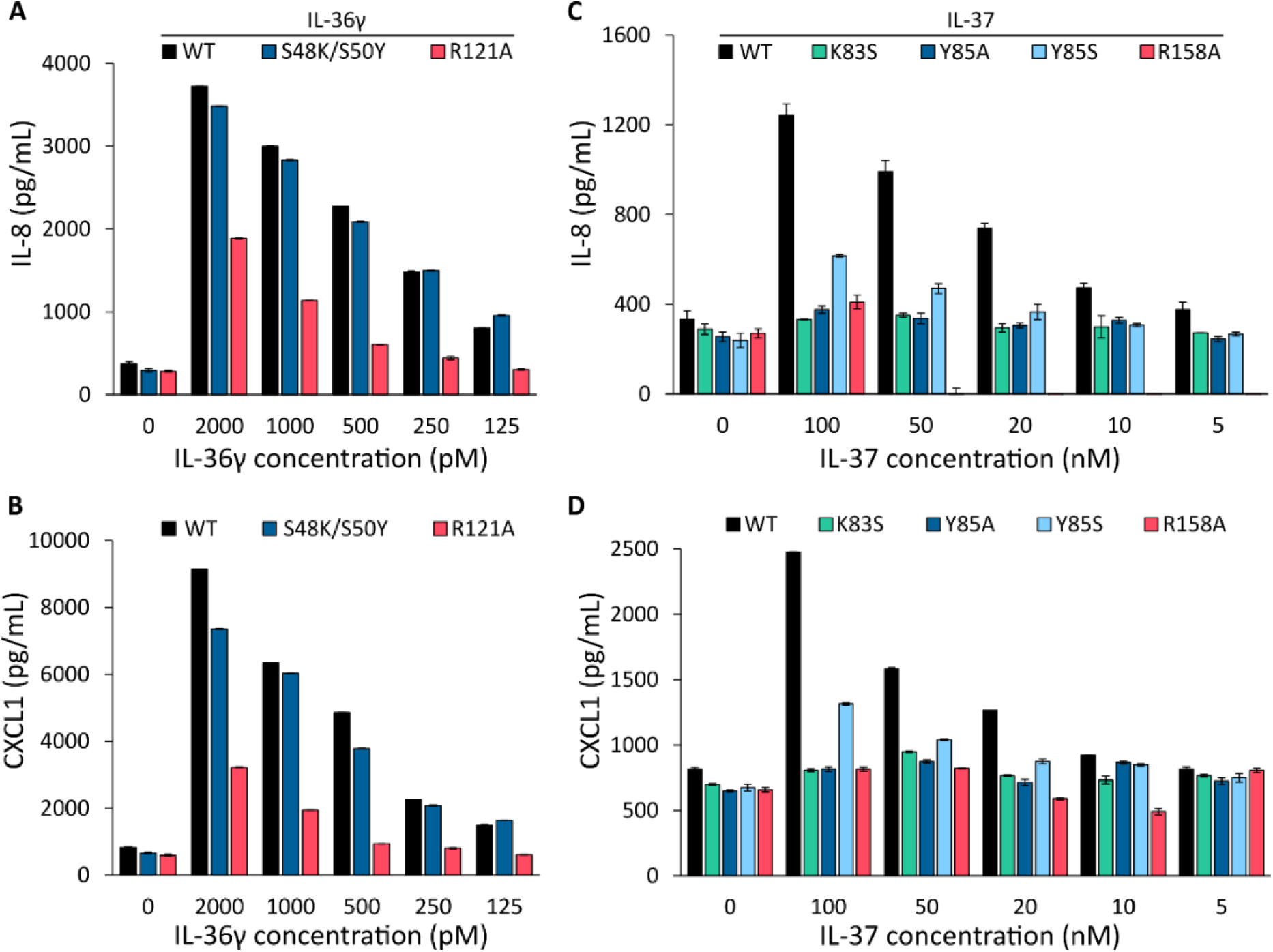
Activity assays of IL-36γ and IL-37 mutants show differential importance of key residues. (**A-D**) HaCaT cells were incubated overnight at 37°C with mature IL-36γ (S18) (A-B), IL-37 (K53) (C-D) and mutants thereof, as indicated. The IL-8 (A, C) and CXCL1 (B, D) concentrations in culture supernatants were determined by ELISA. A representative from three biological replicates is shown. Error bars are from three technical replicates.

Collectively, these results revealed that mutations at the dimerization interface of IL-37 markedly impaired both receptor binding and downstream pro-inflammatory activity, indicating overlapping interfaces. In addition, the conserved arginine residue at Site 2 in the IL-36R interface was shown to be crucial for cytokine-receptor interactions. While the impact of these mutations varied between IL-36γ and IL-37, they underscore the importance of these conserved sites in mediating cytokine function.

## DISCUSSION

Despite the extensive immunological profile of IL-36R signaling in physiology and disease, including the therapeutic targeting of IL-36R by the approved monoclonal antibody spesolimab, the field has been limited by a paucity of structural insights about IL-36R’s signaling complexes. Here, we have abridged this gap in knowledge and have provided structural insights into how IL-36γ mediates a signaling complex comprising IL-36R and the shared receptor IL-1RAcP. Furthermore, we have characterized the interaction of IL-36R with IL-36γ and the recently identified agonist IL-37 and have provided insights into the functional duality of these interactions in a pro-inflammatory context. Our findings set the stage for understanding IL-36R signaling pleiotropy and redundancy, and will help to steer approaches for the therapeutic targeting of IL-36R.

IL-36 cytokines activate NF-κB and MAPK signaling pathways through a shared receptor complex composed of IL-36R and IL-1RAcP to induce cytokine and chemokine production from target cells (*4*). Recently, IL-37 has been identified as a new ligand for IL-36R with a pro-inflammatory signaling signature. In this study, we employed an integrative approach to elucidate the structural and mechanistic basis of the assembly and activation of the pro-inflammatory IL-36 receptor complex mediated by IL-36γ and IL-37 (**Fig. 8**).

**Fig. 8.**
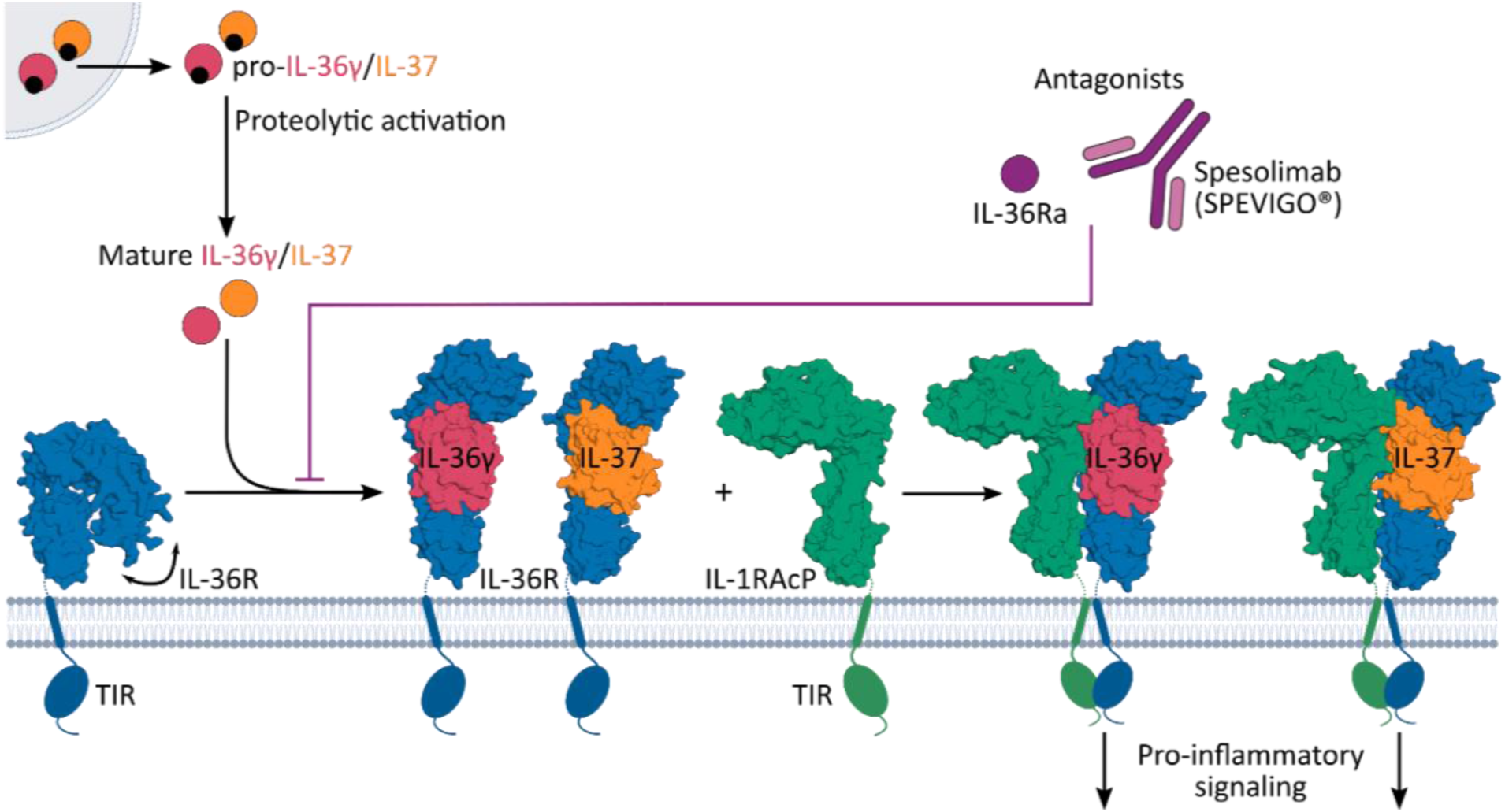
Recapitulation of the structural and mechanistic insights into the assembly and antagonism of the pro-inflammatory complexes. Extracellular proteases released from myeloid cells, e.g., cathepsin G, cathepsin S and elastase, process pro-IL36γ and pro-IL-37 to their mature form. The mature form interacts with IL-36R to induce recruitment of IL-1RAcP and subsequent pro-inflammatory signaling. Reduction of pro-inflammatory signaling occurs upon binding of the naturally occurring IL-36 receptor antagonist (IL-36Ra) or therapeutics, such as spesolimab, to prevent cytokine engagement. The closed conformation of IL-36R on the left represents the AlphaFold Database model. Models with IL-36γ are taken from our cryo-EM structure, and models with IL-37 were predicted using AlphaFold3.

As seen in other IL-1 family receptor systems, IL-36γ initially binds to IL-36R to create a combined interface for IL-1RAcP recruitment, which we confirmed via ITC experiments. Surprisingly, the affinity of the IL-36γ:IL-36R binary complex was relatively low but allowed IL-1RAcP recruitment to create a high affinity, potent pro-inflammatory cytokine-receptor complex (**Fig. 1, A, B, D and E**). The latter was previously assessed only in the context of a heterodimeric Fc fusion of both IL-36R and IL-1RAcP, yielding an up to 70-fold higher affinity, possibly due to avidity effects (*30*, *35*). IL-1RAcP, a shared co-receptor within the IL-1 family, is differentially recruited by distinct cytokine-receptor complexes. For example, IL-1RAcP is primarily engaged by the cytokine interface in the IL-1β complex, whereas the ST2 interface primarily drives IL-1RAcP recruitment in the IL-33 complex (*39*). Here, we present the cryo-EM structure of the IL-36γ ternary complex at a nominal resolution of 3.27 Å. Comparative analysis of the interaction surface with IL-1RAcP created by the IL-36γ:IL-36R, IL-1β:IL-1R1 and IL-33:ST2 complexes revealed that IL-36γ and IL-36R form a less extensive interface with IL-1RAcP, leading to an at least a threefold reduction in affinity for IL-1RAcP (*K*_D_ = 18 ± 4 nM) compared to the affinity of IL-1β:IL-1R1 (*K*_D_ = 0.9 ± 0.2 nM) or IL-33:ST2 (*K*_D_ = 6.6 ± 3.6 nM) for IL-1RAcP (*39*).

In our cryo-EM structure, the IL-36R D3 domain exhibited lower resolution due to its inherent flexibility, as supported by local resolution estimation. This flexibility is consistent with the presence of a linker connecting the D1D2 module to D3 in IL-36R. Previous small-angle X-ray scattering (SAXS) studies have shown similar behavior inST2, which can adopt various conformations distinct from its IL-33-bound form (*29*). Our ITC and SEC experiments revealed that a fraction of IL-36R is unable to interact with the cytokine, indicating the presence of an alternative, potentially inactive conformation. Interestingly, the predicted structure of IL-36R in the AlphaFold database shows a collapsed form where the third domain is folded inward, in a conformation that occludes the cytokine binding interface. A similar conformation was observed for IL-1R1 in complex with the peptide antagonist AF10847, though the physiological relevance remains unclear (*46*).

Since their discovery, IL-36 cytokines have been associated with various inflammatory diseases, including psoriasis, systemic lupus erythematosus (SLE), rheumatoid arthritis (RA), and inflammatory bowel disease (IBD) (*47*, *48*). During homeostasis, IL-36 agonists are regulated by IL-36Ra, their natural antagonist, to prevent hyperinflammation (*5*). However, mutations in *IL36RN*, the gene encoding for IL-36Ra, have been linked to GPP (*9*, *41*, *49*). Spesolimab, an FDA-approved IL-36R-targeting antibody, is used to treat GPP in adults and pediatric patients aged 12 and above. However, the mechanism of action remained unclear (*26*). The overlay of the crystal structure of the spesolimab Fab fragment in complex with IL-36R and the cryo-EM structure of the IL-36γ ternary complex revealed a clash between the D1-D2 loop of IL-36R and the β3-β4 loop of IL-36γ, induced by binding of the Fab fragment. This suggests that spesolimab acts as an allosteric antagonist to inhibit IL-36R driven signaling.

Notably, Ser48 and Ser50 are located in the β3-β4 loop of IL-36γ. Mutating these residues in IL-36γ to the corresponding residues in IL-37, which has a higher affinity for IL-36R, resulted in a slight increase in affinity and similar pro-inflammatory activity in HaCaT cells (Fig. 5B and Fig. 7, A and B). Conversely, mutations of Lys83 and Tyr85 in IL-37 to serine, as in IL-36γ (K83S, Y85S), or to alanine (Y85A) (*42*, *44*) underscored the significance of this loop. Although only the Y85A mutation had a major effect on affinity, both Y85A and Y85S profoundly affected pro-inflammatory activity (**Fig. 5, E and F, and Fig. 7, C and D**). Mutation of Lys83 to serine abolished the detection of an interaction by ITC and eliminated pro-inflammatory activity above the threshold (**Fig. 5D and Fig. 7, C and D**). These findings highlight the critical role of the IL-36γ/IL-37 β3-β4 loop in IL-36R binding, suggesting that the clash with the D1-D2 loop may block cytokine binding, explaining the mechanism of action of spesolimab. Nonetheless, further investigation using cytokine-antibody competition assays are needed to confirm whether spesolimab binding prevents IL-36 or IL-37 cytokine engagement. Given the redundancy and pleiotropy of IL-1 family cytokines and receptors, targeting IL-1RAcP could offer broader therapeutic effects across all cytokines that utilize IL-1RAcP. Two anti-IL-1RAcP monoclonal antibodies, CAN10 and 3G5, have been shown to block IL-1β, IL-33 and IL-36 signaling (*50*). Despite the differential recruitment of IL-1RAcP, several hotspot residues on IL-1RAcP are consistently engaged. The mechanism of inhibition in terms of IL-36γ can now be discussed in the light of our structural data. The epitope of CAN10 is located on the c2d2 loop of IL-1RAcP D2, where Met179, Asn186, Phe187, Asn188 and Ile191 also form part of the interface with IL-36γ:IL-36R (**Table S3**). 3G5 interacts with D3 of IL-1RAcP, disrupting interface residues Tyr269 and Arg306. Further biophysical and structural characterization of IL-37 complexes is needed to evaluate the impact of these antibodies on IL-37 signaling.

IL-37 exists as a dimer in solution with reported dissociation constants ranging from 5 to 294 nM (*21*, *42*). According to the AlphaFold3 model, the dimer interface overlaps with the binding interface with IL-36R. Since for the ITC experiments performed herein IL-37 concentrations in the high micromolar range were used, IL-37 would be expected to be present as a dimer. Because the IL-37 dimer must dissociate before binding to IL-36R, it is possible that the dissociation constant we report for the IL-37:IL-36R interaction determined by ITC represents an apparent value. Titration of dimeric IL-37 into buffer (HBS) showed neglectable heat absorption (Fig. S7E). However, direct measurement of the heat of dimer dissociation is challenging, as the signals remain below background at the low micromolar to nanomolar concentrations required to dissociate the dimer. Additional experiments are required to understand if and how IL-37 dimerization limits its association with the receptor.

The Y85A mutation in IL-37 resulted in a marked decrease in both affinity and pro-inflammatory activity. Interestingly, previous studies have described IL-37 as anti-inflammatory cytokine, with this activity being enhanced in the monomeric Y85A variant (*42*). The second subunit of the IL-37 dimer would clash with IL-36R D1 and D2, suggesting a monomeric variant could be more active (*42*, *44*). We showed that the K83S, Y85S and Y85A mutants are monomeric. However, the β3-β4 loop, which contains these residues, is involved in both the dimer and IL-36R D2 interface, with mutated variants leading to greatly reduced affinity towards IL-36R and pro-inflammatory activity in HaCaT cells. For its previously reported anti-inflammatory effects, IL-37 has been suggested to utilize IL-18R1 and SIGIRR (*19*). It is possible that the interface with these receptors is not affected by the Tyr85 mutation in IL-37.

The mechanism by which IL-37 exhibits both pro- and anti-inflammatory activity remains unclear. Akin to other IL-1 cytokines, IL-37 is produced as an inactive precursor with an N-terminal pro-domain that requires processing to achieve full activity. In this study, we confirmed the pro-inflammatory activity of Lys53-IL-37, while Val46-IL-37 and His47-IL-37 did not induce IL-8 production in HaCaT cells (*18–20*). Val46-IL-37 was identified through N-terminal sequencing of IL-37 released from HEK293 cells post-transfection (*18*). Although the protease responsible for processing IL-37 before Val46 is unknown, elastase is known to cleave after Val46, yielding His47-IL-37 (*20*). Despite this, Val46-IL-37 has been extensively used over the past two decades, in addition to pro-IL-37 transfection in a variety of cell lines or transgenically overexpressing mouse models, without confirmation that the necessary proteases are present or that effective proteolytic processing occurs (*18*, *19*, *42*). IL-36 family cytokines are expressed with very short N-terminal propeptides (ranging from 4 to 17 residues), however, incorrect processing by even a single amino acid in any of these cytokines leads to a 1,000- to 10,000-fold reduction in activity, comparable to the inactive full-length pro-form (*13*). The critical importance of correct pro-domain processing is further highlighted by the structure of pro-IL-18, which reveals a conformation markedly different from mature IL-18 (*51–53*). Alternative processing of IL-37 could lead to different conformations, engaging different receptor complexes and producing varied signaling outcomes (*53*).

Collectively, the structural and mechanistic insights provided here will guide further investigation of the pro-inflammatory activation of IL-36R by IL-36γ and IL-37. These findings will not only enhance our understanding of IL-36R sharing, but also open new avenues for therapeutic intervention, considering the emerging pro-inflammatory profile of IL-37. Finally, further research will be needed to investigate the relevance of the conformational plasticity of IL-36R in engaging different cytokines agonists to elicit finetuned signaling outputs depending on context.

## MATERIALS AND METHODS

### Study Design

The purpose of this study was twofold: (1) to generate structural and mechanistic insights into how human IL-36R, a clinical target in acute skin inflammation, nucleates a signaling complex mediated by its best understood cytokine ligand IL-36γ, and (2) to obtain a framework for understanding the structure-function landscape of IL-37, which was recently proposed to signal via IL-36R with a pro-inflammatory signature.

We first produced a plethora of recombinant proteins in bacterial and mammalian cells and established purification protocols to obtain highly pure and monodisperse proteins and protein complexes. We have used these recombinant proteins to carry out interaction studies by isothermal titration calorimetry (ITC) towards quantifying binding affinities, stoichiometries and thermodynamic binding parameters. We further employed these recombinant proteins in cellular assays based on HaCaT cells to evaluate pro-inflammatory signaling mediated by IL-36γ and IL-37. We reconstituted biochemically the human IL-36γ:IL-36R:IL-1RAcP complex and used a purified complex to determine its structure by single-particle cryo-electron microscopy. Structural models involving IL-37 were obtained by AI-based structure predictions. We used the structural models derived by experiment and computation in conjunction with protein sequence information including from orthologous proteins to sketch the structure-function landscape of cytokine-mediated activation of IL-36R. This information formed the basis for structure-guided mutagenesis of IL-36γ and IL-37 to interrogate their utilization of IL-36R and to provide a framework for the duality of IL-36R.

### Recombinant protein expression constructs

Human cDNA sequences for the cytokines were codon-optimized for protein expression in *Escherichia coli* and purchased from IDT (Integrated DNA Technologies) as geneblocks. The sequences were cloned into the pET15b vector (Novagen, Merck) in frame with an N-terminal caspase-3 cleavable hexahistidine (His_6_)-tag using a traditional restriction-ligation approach. Sequences encoding the mature version of IL-36γ (UniProt ID Q9NZH8) comprising amino acids 18-169, and of IL-36Ra (UniProt ID Q9UBH0) comprising amino acids 2-155, were used. In the case of IL-37 (isoform b, UniProt ID Q9NZH6), different mature constructs were designed, including IL-37_V46_ (residues 46-218), IL-37_H47_ (residues 47-218) and IL-37_K53_ (residues 53-218). In addition, constructs carrying mutations were designed, including IL-36γ_S48K/S50Y_, IL-36γ_R121A_, IL-37_K83S_, IL-37_Y85A_ and IL-37_Y85S_. All IL-37 mutants have K53 as starting residue at the N-terminus.

Sequences encoding the extracellular part of the human receptors were codon-optimized for protein expression in mammalian HEK293 cells and purchased from IDT (Integrated DNA Technologies) as geneblocks. Sequences of the extracellular domains containing the native signal peptide were used for IL-36R (UniProt ID Q9HB29, residues 1-335) and IL-1RAcP (UniProt ID Q9NPH3, residues 1-367). To improve the stability of IL-36R, two cysteine to serine mutations were introduced (C154S, C262S). Since SIGIRR is a single pass Type III membrane protein, the signal peptide, contained in the transmembrane region, is not part of the extracellular sequence. Therefore, the sequence of SIGIRR (UniProt ID Q6IA17, residues 1-118) was preceded by the chicken RTPµ-like signal peptide sequence (*54*). The extracellular receptor fragments were cloned into the pHLSec vector (Addgene plasmid #99845, (*54*)) in frame with a C-terminal caspase-3 cleavable His_6_-tag using a traditional restriction-ligation approach. All constructs were validated by Sanger sequencing by Eurofins Genomics.

### Protein expression in *E. coli* and purification

BL21(DE3) *E. coli* transformed with a pET15b plasmid expressing N-terminally His_6_-tagged cytokine (IL-36γ, IL-36Ra, IL-37_V46_, IL-37_H47_, IL-37_K53_, IL-36γ_S48K+S50Y_, IL-36γ_R121A_, IL-37_K83S_, IL-37_Y85A_ or IL-37_Y85S_) were grown at 37°C in LB medium supplemented with carbenicillin (100 µg ml^-1^) as selection marker for the pET15b plasmid. When the optical density at 600 nm reached 0.6, the expression of the cytokines was induced by addition of isopropyl β-D-1-thiogalactopyranoside (IPTG) at a final concentration of 0.1 mM. The culture was incubated overnight (21h) at 20°C, harvested the next day by centrifugation (4000g for 15 min at 4°C) and the cellular pellet was stored at −80°C.

The bacterial pellet was thawed and resuspended in HEPES-buffered saline (HBS, 20 mM HEPES, 150 mM NaCl, pH 7.4). The cells were lysed by sonication with a Qsonica macrotip sonicator (4 min on-time, pulse on 30”, pulse off 30”, amplitude 70%) while cooled on ice. The cell debris was removed by centrifugation (20,000g for 1h at 4°C) and the clarified lysate was filtered using a 0.22 µm Millipore Steritop Sterile Vacuum Bottle-Top Filter (Merck). The filtered lysate was loaded onto a 5 ml HisTrap FF column (Cytiva) and the His-tagged cytokine was eluted using HBS supplemented with 250 mM imidazole, followed by desalting using a HiPrep 26/10 Desalting column (Cytiva) to remove imidazole. The His-tag was removed by incubation with caspase 3 (produced in-house) at 20°C overnight, after which the cleaved protein was purified by loading onto a 5 ml HisTrap FF column (Cytiva) and collecting the flowthrough. As a polishing step, the sample was concentrated and injected onto a Superdex 75 increase 10/300 GL column (Cytiva) and fractions containing the cytokine were pooled, flash frozen and stored at −80°C. The purity of the cytokines was evaluated by SDS-PAGE analysis.

### Protein expression in HEK293 cells and purification from conditioned medium

The extracellular domains (ECD) of the receptors (IL-36R, IL-1RAcP) were produced in suspension adapted HEK293S cells (kindly provided by Prof. Nico Callewaert, VIB Center for Medical Biotechnology, Belgium). Cells were maintained in a mixture of 50% Freestyle 293 Expression Medium (Gibco) and 50% Ex-Cell 293 Serum-Free Medium (SAFC, Merck), and diluted to 1 x 10^6^ cells ml^-1^ the day before transfection. Transient transfection was performed using linear polyethylenimine 25 kDa (Poly-sciences) as transfection reagent. One day post-transfection, valproic acid was added to a final concentration of 1.5 mM, and glucose to a final concentration of 5 mg ml^-1^. Five days post-transfection, conditioned medium was clarified by centrifugation at 1,000g for 15 min and filtered through a 0.22 µm filter prior to chromatographic purification steps. The C-terminally His-tagged receptors were captured onto a 5 ml cOmplete His-tag Purification Column (Roche). The protein was eluted using HBS supplemented with 250 mM imidazole, which was removed using a HiPrep 26/10 Desalting column (Cytiva). The His-tag was removed by incubation with caspase 3 (produced in-house) at 20°C overnight, after which the cleaved protein was purified by loading onto a 5 ml HisTrap FF column (Cytiva) and collecting the flowthrough. As a polishing step, the sample was concentrated and injected onto a Superdex 200 increase 10/300 GL column (Cytiva) and fractions containing the receptor were pooled, flash frozen and stored at −80°C. The purity of the receptors was evaluated on SDS-PAGE.

### SEC-MALLS and mass photometry

For each sample, 120 µg (80 µl at 1.5 mg ml^-1^) was injected onto a Superdex 200 increase 10/300 GL column (Cytiva) pre-equilibrated in HBS, connected to an Agilent HPLC system with an autosampler. The column was coupled in line to an ultraviolet detector (Agilent), a DAWN8 (Wyatt) MALLS detector and an Optilab (Wyatt) refractometer. The data was analyzed using the ASTRA8.2 software (Wyatt). Glycosylated receptors were subjected to protein conjugate analysis. Refractive index increment values (dn/dc) of 0.185 ml g^-1^ and 0.165 ml g^-1^ were used for protein and glycan analysis, respectively. Bovine serum albumin (BSA, Pierce) was used as a standard to correct for band broadening.

Mass photometry experiments were performed on a Refeyn TwoMP mass photometer (Refeyn). Calibration of the instrument was performed with β-amylase (BAM) and IgG at a concentration of 5 nM and 10 nM, respectively. Samples were diluted to 10 nM in PBS prior to measurement. Three technical replicates were overlayed into one graph. Acquisitions were performed in AcquireMP 2023 (Refeyn) and analysis in DiscoveryMP 2023 (Refeyn).

### IL-36γ:IL-36R:IL-1RAcP complex reconstitution and cryo-EM data collection

IL-36γ was incubated with IL-36R and IL-1RAcP in a 1:1:1 molar ratio for 30 minutes before injection onto a Superdex 200 increase 10/300 GL column (Cytiva) to purify the ternary complex. Fractions corresponding to the complex were pooled, concentrated to 4.1 mg ml^-1^, and aliquots were flash-cooled into liquid nitrogen and stored at −80 °C until further use. The presence of all three components was validated by SDS-PAGE. Immediately before application onto glow-discharged Quantifoil^TM^ R 2/1 on 300 copper mesh grids, dodecyl maltoside (DDM) was added to a final concentration of 0.1 mM, resulting in protein sample dilution to 3.3 mg ml^-1^. After application of 4 µl of sample, grids were blotted for 4.5 s under 95% humidity at 22°C and plunged into liquid ethane using a Leica EM GP2 Plunge Freezer. Grid screening was performed on a JEOL JEM-1400plus microscope at the VIB-UGent Bioimaging Core (Ghent, Belgium). Data collection was performed using a 300 kV CryoARM300 microscope (JEOL) equipped with an Omega filter (JEOL) and a K3 direct electron detector (Gatan), operated using SerialEM version 4.1 (*55*). A total of 8,828 movies were collected at a magnification of 60,000x, corresponding to a raw pixel size of 0.74 Å.

### Cryo-EM image processing

Image processing was performed in cryoSPARC v4.2.1 (*56*) (Fig. S3). Movies were motion-corrected and dose weighted using Patch motion correction, and Patch contrast transfer function (CTF) estimation was performed on the corrected micrographs. Initial particle picking was performed using crYOLO (*57*) resulting in 771,846 particles, which were imported into cryoSPARC. The particles were extracted using a box size of 360 pixels and downsampled (2x binned) to a box size of 180 pixels, corresponding to a pixel size of 1.48 Å per pixel. After multiple cycles of 2D classification, particles corresponding to ten representative 2D class averages were selected to train Topaz (*58*). Subsequent particle extraction resulted in 643,451 particles that were used for another round of 2D classification. Selected 2D class averages were used to generate *ab initio* 3D reconstructions, which were heterogeneously refined into five classes. The dominant class was further processed using homogeneous and non-uniform refinement. After another round of heterogeneous, homogeneous and non-uniform refinement, particles were reextracted in a box size of 360 pixels corresponding to a pixel size of 0.74 Å per pixel. A final non-uniform refinement resulted in a final map at a nominal resolution of 3.3 Å based on the 0.143 gold-standard Fourier shell correlation (FSC) criterion (*59*). The cryo-EM map was further sharpened by applying a *B* factor −70 Å^2^ for model building and visualization.

### Cryo-EM model building and refinement

A structural model for the ternary complex consisting of IL-36γ, IL-36R and IL-1RAcP was predicted using AlphaFold-Multimer version 2.2 (*60*). The predicted model was rigid-body fitted in the sharpened map using UCSF ChimeraX (*61*) and subsequently subjected to automatic MD flexible fitting using NAMDINATOR (*62*). The model was iteratively adjusted by several cycles of manual building in Coot (*63*) and real-space refinement in Phenix (*64*) (version 1.20.1-4487) using the sharpened map. Validation of the model was performed using Phenix (*64*) and MolProbity (*65*). Cryo-EM data collection, refinement and validation statistics are summarized in **Table S2**. Visualization of cryo-EM maps and structural models was performed using UCSF ChimeraX (*61*) and PyMol (The PyMOL Molecular Graphics System, Version 2.3.3, Schrödinger, LLC).

### *In silico* structure prediction and bioinformatic analysis

A structural model for the ternary complex consisting of IL-36γ, IL-36R and IL-1RAcP was predicted using AlphaFold-Multimer version 2.2 (*60*) and used to generate a starting model for real-space cryo-EM refinement. In addition, a structural model for IL-37 in complex with IL-36R was predicted by AlphaFold3 using the DeepMind server (*66*) and aligned with the IL-36γ:IL-36R complex of the cryo-EM structure in PyMol (The PyMOL Molecular Graphics System, Version 2.3.3, Schrödinger, LLC).

Multiple sequence alignment was performed using Clustal Omega (*67*) and visualized using ESPript3.0 (*68*). PDBePISA (*40*) was used for protein-protein interaction analysis, and interfacing residues were indicated on the sequence alignment.

### Isothermal titration calorimetry

In the last purification step, all proteins were buffer matched in the same HBS buffer via size exclusion chromatography (SEC). Protein concentrations were determined with the NanoDrop 2000 (Thermo Fisher Scientific) using their corresponding extinction coefficients (absorbance of 1%) as determined with the ProtParam tool (ExPASy) (*69*). Experiments were carried out on the MicroCal PEAQ-ITC instrument. Titrations were preceded by an initial injection of 0.4 µl. The titration consisted either of 12 injections of 3 µl during 6 sec each, or of 19 injections of 2 µl during 4 sec each, with 150 sec spacing between injections. All experiments were carried out at 25 °C. The cell concentration was aimed at 19 µM, while the syringe concentration was aimed at 190 µM. While performing the ternary titrations where IL-1RAcP was titrated into the binary complex of IL-36γ (WT or mutants) and IL-36R, the sample cell of the binary titration was retained albeit after removing the excess overflowing from the sample cell. Throughout the experiment, the sample was stirred at a speed of 750 rpm. Data was initially analyzed using PEAQ-ITC analysis software (version 1.1.0.1262, Malvern) and fit using a ‘one set of sites’ model. To improve the accuracy of the thermodynamic parameters, a global numerical fitting was performed using KinTek Explorer 10 (KinTek Corporation, USA).

### Global numerical analysis of ITC data

The analysis of ITC data was performed by global numerical fitting using KinTek Explorer 10 (KinTek Corporation, USA) (*31*, *32*). The software allows for the input of a given kinetic model via a simple text description, and the program then derives the differential equations needed for numerical integration automatically. The defined kinetic model (Equation 1), used as a base for the data fitting, contained three major steps based on other supporting experiments: (i) presence of two forms of the IL-36R receptor in the solution (inactive IL36R_inact_ vs. active IL36R_act_); (ii) binding of a cytokine (C) to IL36R_act_ to form a binary complex; (iii) additional sequential binding of IL-1RAcP to form a ternary complex. By the definition of the model provided in Equation 1, strict 1:1:1 binding was always assumed during the data fitting, corresponding to fixing the *n*-value to 1 in conventional ITC data fitting. Kinetics of the first step (IL-36R inactive vs. active fraction) was set to 10^6^-fold slower rate constants (10^-4^ s^−1^), followed by extensive pre-equilibration before simulating the cytokine addition. Such arrangement resulted in negligible concentration changes between the two species (inactive vs. active) within the experimental window and mimicked two separate populations of IL-36R without the possibility of re-equilibration upon cytokine binding. Furthermore, forward rate constants for both the binary and the ternary complex formation steps were fixed to 100 s^−1^ to mimic the rapid equilibrium assumption. This allowed reliable and precise estimation of equilibrium affinities (*K*_d,bin_ and *K*_d,ter_ for the binary and the ternary complexes, respectively) without overfitting data with elementary rate constants (*k*_a_, *k*_d_) of each step that were not well-defined by the equilibrium data. When no subsequent binding of IL-1RAcP was observed for a given IL-36R.cytokine binary complex, the rate constant of the ternary complex formation was fixed to 0 s^−1^ to exclude this step from the defined mechanism and analysis.

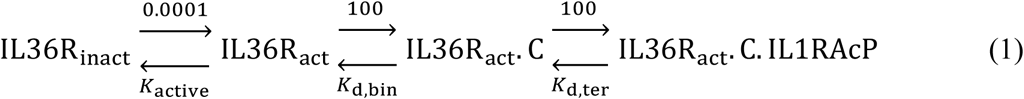

Raw ITC data were integrated to obtain dependence of the cumulative heat released/consumed with increasing concentration of the ligand injected into the cell and the data were fit globally into the defined kinetic model (Equation 1) using Equation 2 as a defined observable. Q corresponds to the released/consumed heat, Δ*H*_bin_ to the enthalpy of the binary complex formation, Δ*H*_ter_ to the enthalpy of the ternary complex formation, [IL36R_act_.C] to the concentration of the receptor.cytokine binary complex, and [IL36R_act_.C.IL1RAcP] to the concentration of the ternary receptor.cytokine.IL1RAcP complex.

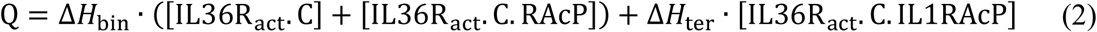

Numerical integration of rate equations searching for a set of kinetic parameters that produce a minimum χ^2^ value was performed using the Bulirsch–Stoer algorithm with adaptive step size. Nonlinear regression to fit the data was based on the Levenberg–Marquardt method. Residuals were normalized by sigma values for each data point. Standard errors were calculated from the covariance matrix during nonlinear regression. A more rigorous analysis of the variation of the kinetic parameters was achieved by performing a confidence contour analysis using FitSpace Explorer (KinTek Corporation, USA)(*33*). In these analyses, the lower and upper limits for each parameter were derived from the confidence contour by setting the χ^2^ threshold at 0.95. The final fit provided a precise and uniquely constrained estimation of enthalpies and dissociation constants for both the binary and the ternary complex, along with the estimation of the inactive portion of the IL-36R receptor.

### Cellular assays

HaCaT cells were maintained in DMEM medium (Gibco) supplemented with 10% fetal calf serum (FCS, Biowest) at 37°C in a humidified atmosphere with 5% CO_2_. For cytokine stimulation, HaCaT cells were plated at 5 x 10^4^ cells per well in 24 well plates in DMEM with 10% FCS supplemented with penicillin/streptomycin (10^6^ units l^-1^ penicillin, 1 g l^-1^ streptomycin, Gibco). Cytokines were added at the concentrations, and cells were incubated overnight at 37 °C. For inhibition assays with IL-36Ra, cells were pretreated with IL-36Ra for 1h at 37 °C, after which 20 nM of IL-36γ or IL-37 variants were added. After cytokine treatment, cell culture supernatants were collected and the concentration of pro-inflammatory cytokines IL-8 and CXCL1 were measured by ELISA (Duoset, R&D). All cytokine assays were carried out using triplicate samples.

### Statistical analysis

ITC data were initially analyzed using the PEAQ-ITC analysis software (version 1.1.0.1262, Malvern) and fit using a ‘one set of sites’ model. Standard deviation values were derived from at least three replicates. To improve the accuracy of the thermodynamic parameters, a global numerical fitting was performed by global numerical fitting using KinTek Explorer 10 (KinTek Corporation, USA) (*31*, *32*). All cellular assays resulting in cytokine release measurements were carried out using triplicate samples.

## Acknowledgements

We thank M. Fislage at the VIB-VUB facility for Biological Electron Cryogenic Microscopy for assistance in cryo-EM data collection, technical support and infrastructural access. We thank K. Balcaen and J. Haustraete at the VIB Protein Core facility (Ghent, Belgium) for access to the SEC-MALLS instrument and technical support.

## Author contributions

Conceptualization: SNS

Methodology: JA, DMC, SNS

Investigation: JA, DMC

Visualization: JA, SNS

Formal analysis: JA, MT, JF, DMC, SNS

Funding acquisition: JA, SNS

Supervision: SNS

Writing – original draft: JA, SNS

Writing – review & editing: JA, MT, JF, DMC, SNS

## Data and materials availability

Cryo-EM map and structure have been deposited in the Electron Microscopy Data Bank (EMDB) and the Protein Data Bank (PDB) under accession codes, EMD-52665 and 9I7X respectively.

## SUPPLEMENTARY MATERIALS

### Supplementary Figures

**Fig. S1:**
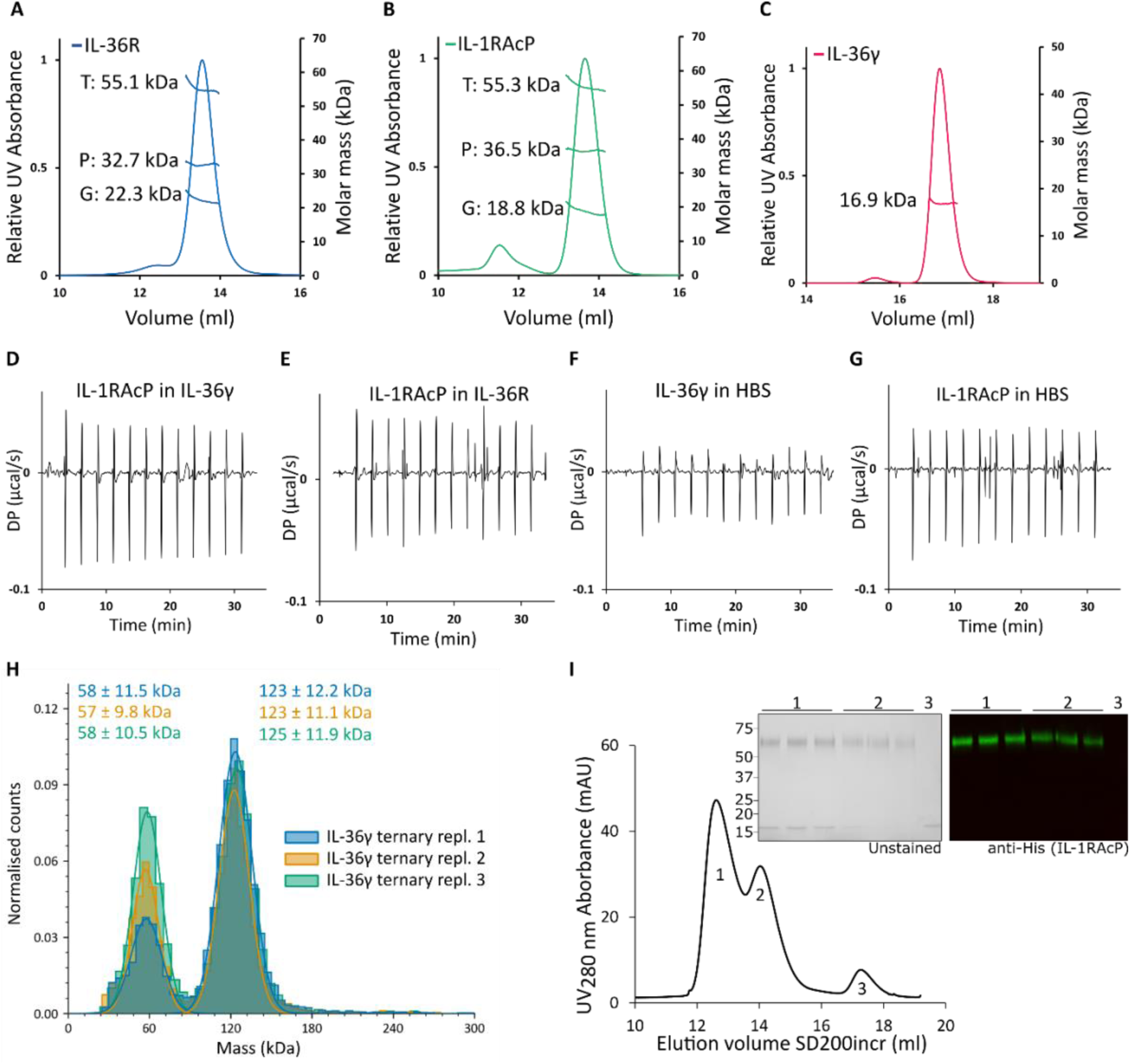
Biochemical and biophysical characterization of IL-36R, IL-1RAcP, IL-36γ and complexes thereof. (**A-C**) SEC-MALLS analysis of IL-36R, IL-1RAcP and IL-36γ, with theoretical masses of 35.9, 39.8 and 17 kDa, respectively. Number of samples analysed: *n* = 1. (A-B) Protein-conjugate analysis of the extracellular domains of IL-36R (A) and IL-1RAcP (B). The SEC-elution profile is plotted as the relative ultraviolet absorbance at 280 nm (left Y-axis) in function of elution volume. The reported total (T), protein (P) and glycan (G) molecular mass (right Y-axis) as determined by MALLS are the average molecular mass throughout the elution peak. Samples were injected onto a Superdex 200 Increase 10/300 GL column (Cytiva). (**D-G**) ITC control experiments between IL-1RAcP and IL-36γ (D) or IL-36R (E) separately. Heat of dilution measurements of IL-36γ (F) and IL-1RAcP (G) in HBS buffer. Abbreviations: DP, differential electrical power. (**H**) Mass photometry analysis of the IL-36γ ternary complex at 10 nM. (**I**) SEC-elution profile and corresponding SDS-PAGE and Western blot analysis of the reconstitution of the IL-36γ ternary complex with C-terminally His-tagged IL-1RAcP. Sample was injected onto a Superdex 200 Increase 10/300 GL (SD200incr) column (Cytiva).

**Fig. S2:**
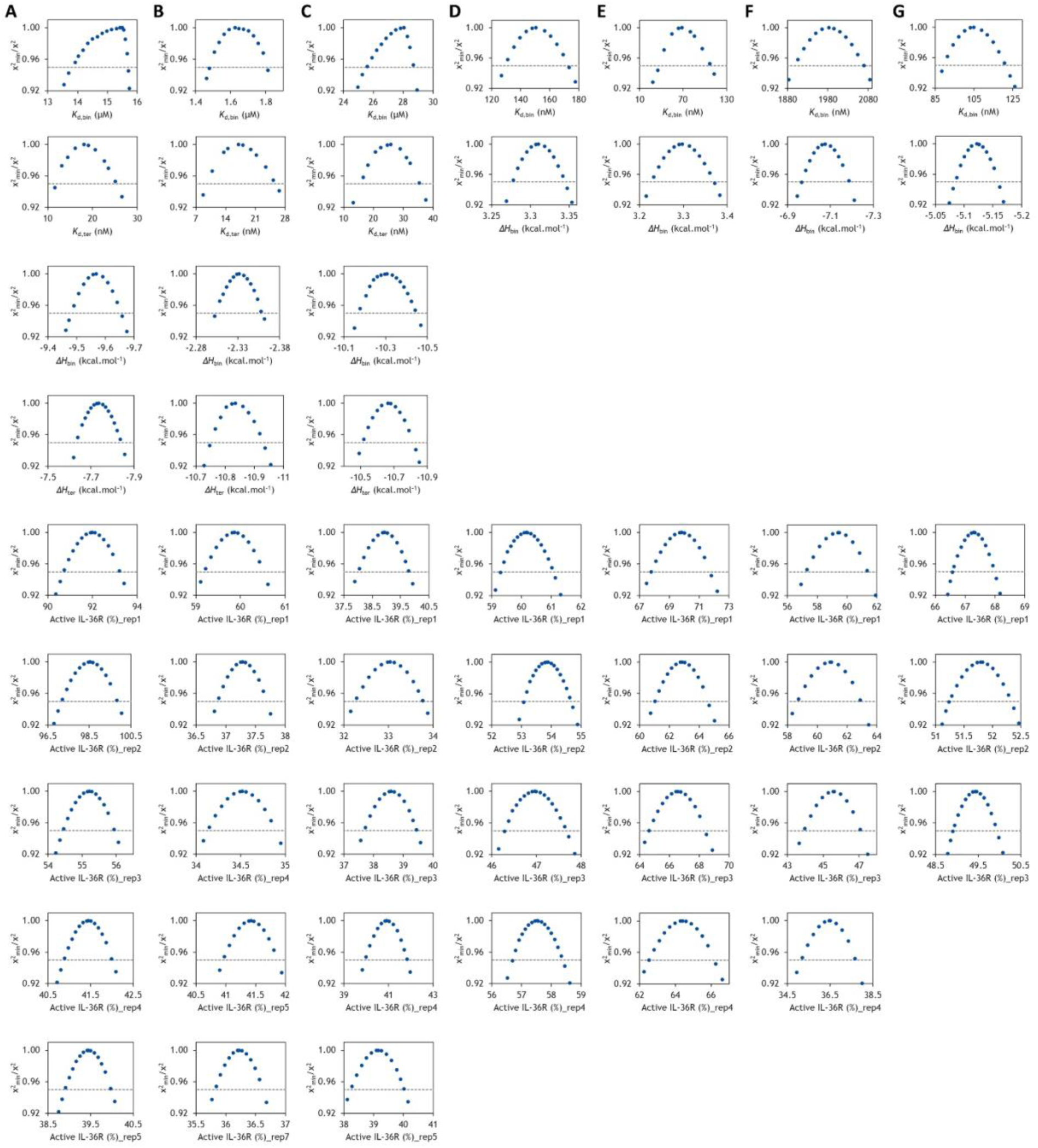
Confidence contour analysis of parameters from global numerical fitting shows that the ITC data is well constrained and robustly analyzed. The statistics are provided for all the tested cytokine variants IL-36γ (**A**), IL-36γ_S48K/S50Y_ (**B**), IL-36γ_R121A_ (**C**), IL-36Ra (**D**), IL-37 (**E**), IL-37_Y85A_ (**F**), and IL-37_Y85S_ (**G**). The grey dashed lines represent the χ^2^ threshold of 0.95. Confidence contour analysis was based on the data fitting shown in Fig. 1, fig. 4, fig. 5 and fig. 6. *K*_d,bin_ and *K*_d,ter_ correspond to dissociation constants of the binary cytokine:IL-36R receptor complex and the ternary cytokine:IL-36R:IL-1RAcP complex, respectively. ΔH_bin_ and ΔH_ter_ correspond to interaction enthalpies of the complex formation for the binary cytokine:IL-36R complex and the ternary cytokine:IL-36R:IL-1RAcP complex, respectively.

**Fig. S3:**
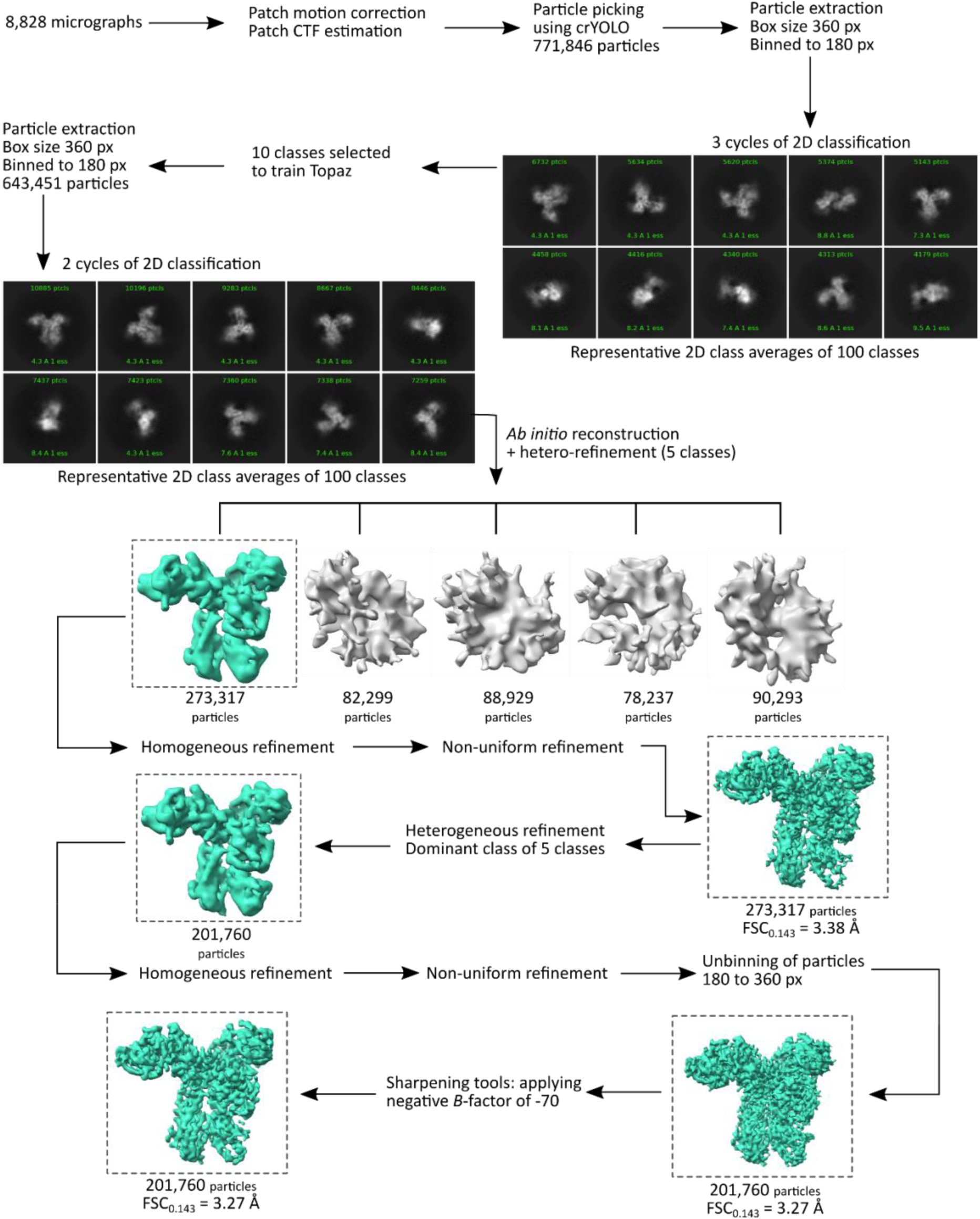
Cryo-EM data processing workflow in cryoSPARC v4.2.1. The estimated resolution at FSC = 0.143 is reported. Abbreviations: CTF, Contrast Transfer Function; FSC, Fourier shell correlation.

**Fig. S4:**
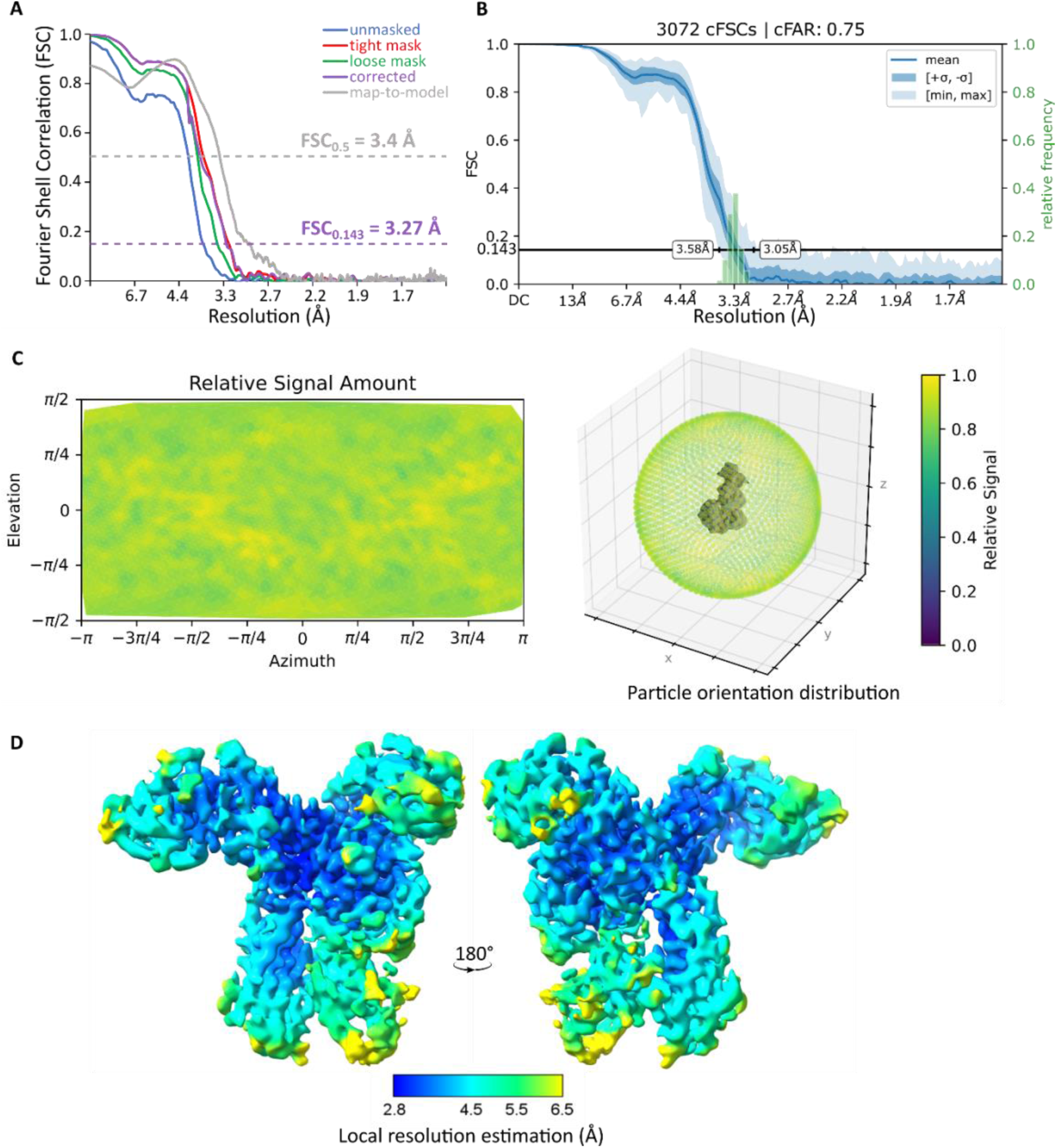
Gold-standard Fourier Shell Correlation (FSC) curves, particle orientation and local resolution estimation. (**A**) Gold-standard FSC curves of the resulting 3D map, calculated after applying either no mask (blue), a loose mask (green), or a tight mask (red) to both half maps. The corrected FSC (purple) is calculated using the tight mask with correction by noise substitution (*70*). The estimated resolution at FSC = 0.143 is shown for the corrected FSC curve (dotted purple line). A map-to-model FSC curve, calculated using Phenix, is shown in grey, along with the estimated resolution at FSC = 0.5 (dotted grey line). (**B**) 3D FSC plot as calculated by the Orientation Diagnostics job in cryoSPARC v4.5.1. (**C**) Particle orientation distribution shown by the Azimuth plot and 3D projection on particle. (**D**) Sharpened map colored according to local resolution.

**Fig. S5:**
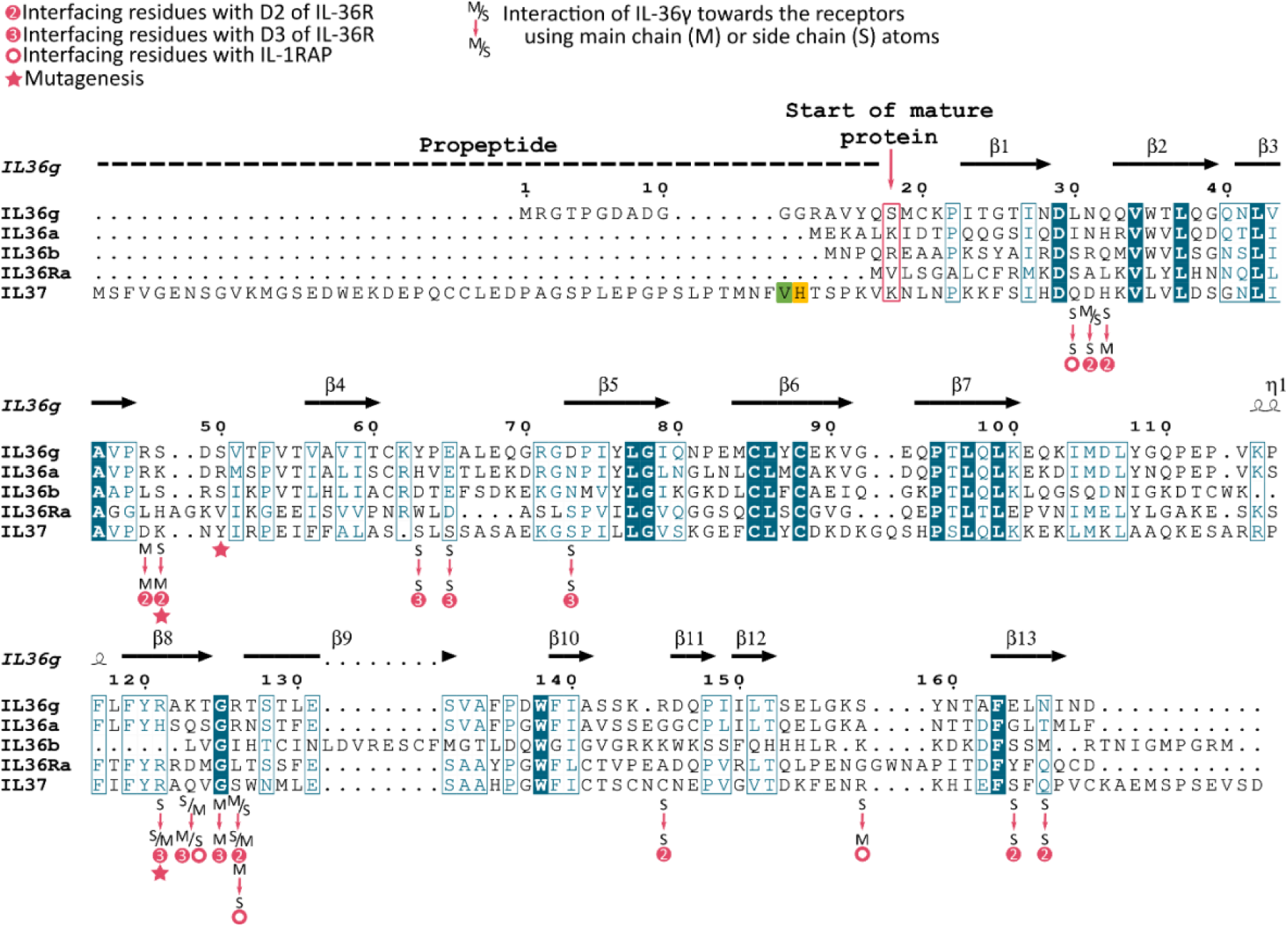
Sequence alignment of IL-36 agonists, IL-36 receptor antagonist (IL-36Ra) and IL-37. The multiple sequence alignment was conducted using Clustal Omega and visualized using ESPript 3.0. Secondary structure elements as found in our cryo-EM structure are indicated above the alignment. Residues of IL-36γ interacting with either IL-36R or IL-1RAcP are indicated according to the legend, also specifying involvement of main chain (M) or side chain (S), related to Table S3. Mutated residues in IL-36γ in this study are indicated with a pink star. Numbering is according to the top sequence. Alternative starting residues at the N-terminus for IL-37 are colored in green and yellow, related to Fig. S7A.

**Fig. S6:**
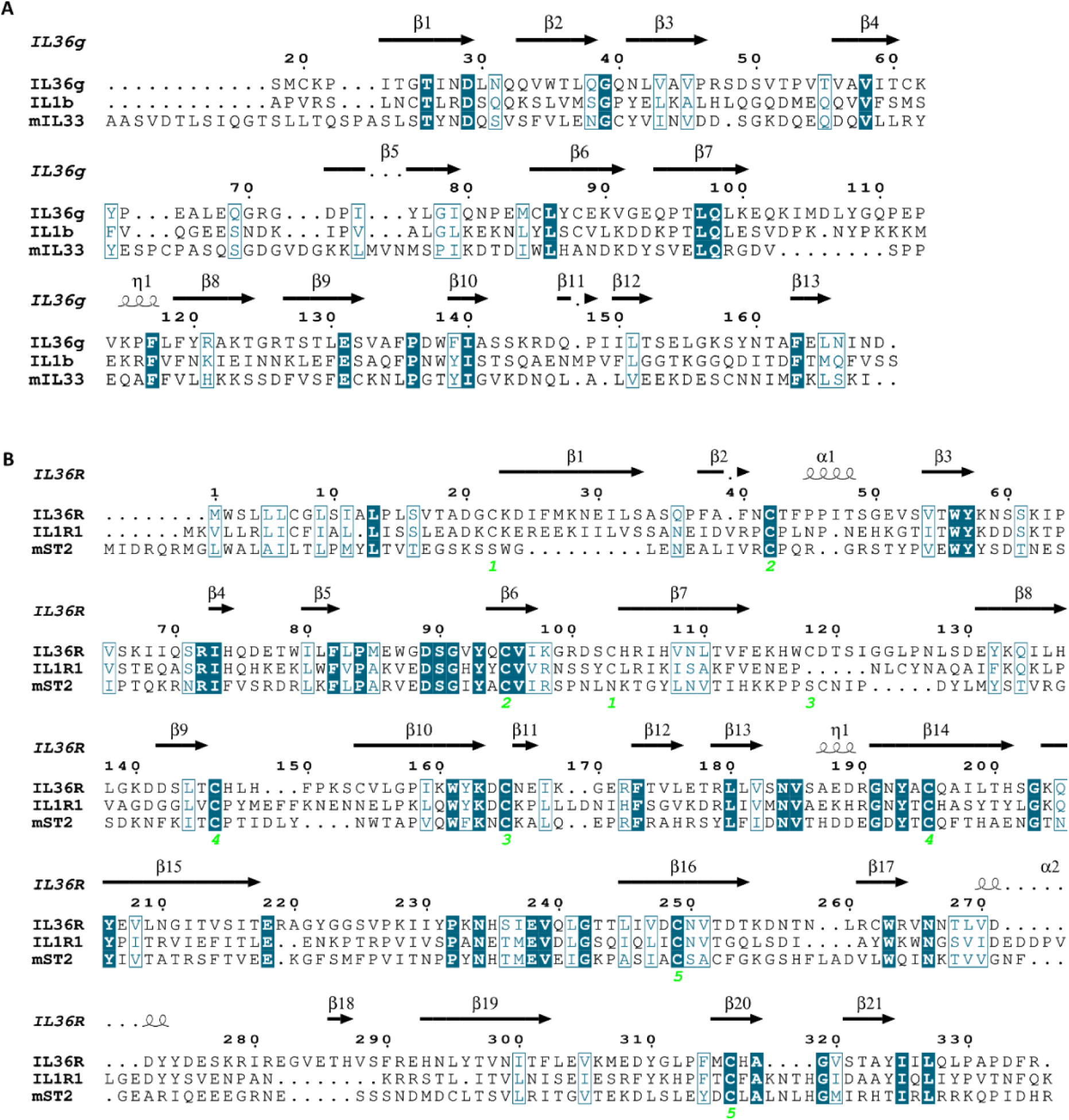
Sequence alignments of selected IL-1 cytokines and receptors. (**A**) Multiple sequence alignment of human IL-36γ, human IL-1β and mouse IL-33, related to Fig. 2C and Table S4. (**B**) Multiple sequence alignment of human IL-36R, human IL-1R1 and mouse ST2, related to Fig. 2D and Table S5. Disulfide bridges, based on the cryo-EM structure of the IL-36γ ternary complex, are indicated by the green numbers underneath the alignment. Multiple sequence alignments were conducted using Clustal Omega and visualized using ESPript 3.0. Secondary structure elements as found in our cryo-EM structure are indicated above the alignment. Numbering is according to the top sequence.

**Fig. S7:**
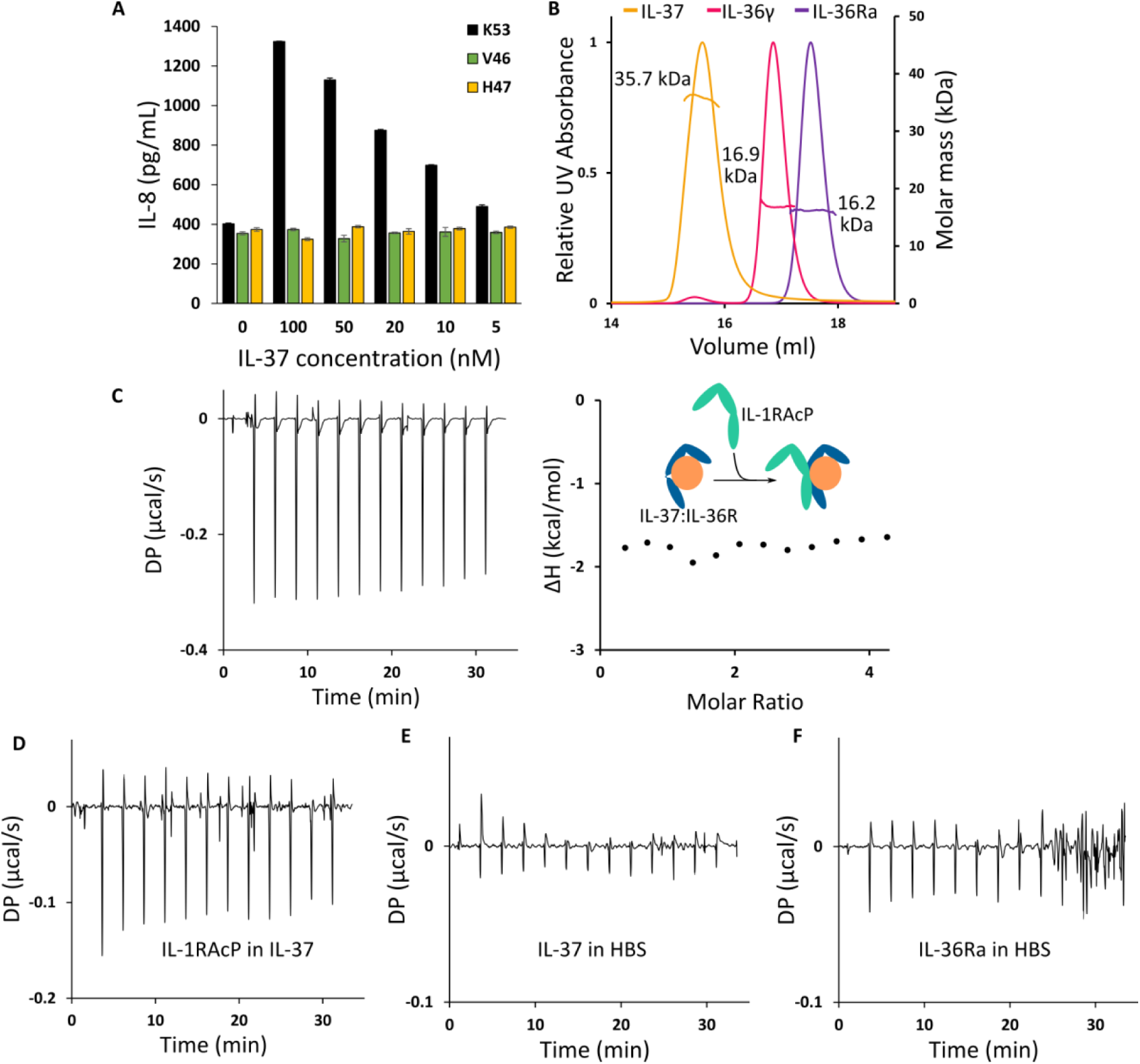
Control experiments for the activity of recombinant variants of IL-37 and biochemical/biophysical characterization of diverse recombinant protein tools. (**A**) HaCaT cells were incubated overnight at 37°C with mature IL-37 starting at Val46, His47 or Lys53. IL-8 concentrations in culture supernatants were determined by ELISA. (**B**) SEC-MALLS analysis of IL-37, IL-36γ and IL-36Ra. The SEC-elution profile is plotted as the relative ultraviolet absorbance at 280 nm (left Y-axis) in function of elution volume. The reported molecular mass (right Y-axis) as determined by MALLS is the average molecular mass throughout the elution peak. The theoretical masses are 18.5 kDa, 17 kDa and 16.8 kDa for monomeric IL-37, IL-36γ and IL-36Ra, respectively. Samples were injected onto a Superdex 200 Increase 10/300 GL (SD200incr) column (Cytiva). Number of samples analysed: *n* = 1. (**C**) Isothermal titration calorimetry experiment of the addition of IL-1RAcP (427 µM) to the IL-37:IL-36R complex (19 µM). (**D-F**) Isothermal titration calorimetry control experiments to test the interaction between IL-1RAcP and IL-37 (D) or the heat of dilution of IL-37 (E) or IL-36Ra (F) in HBS buffer. Abbreviations: DP, differential electrical power.

**Fig. S8:**
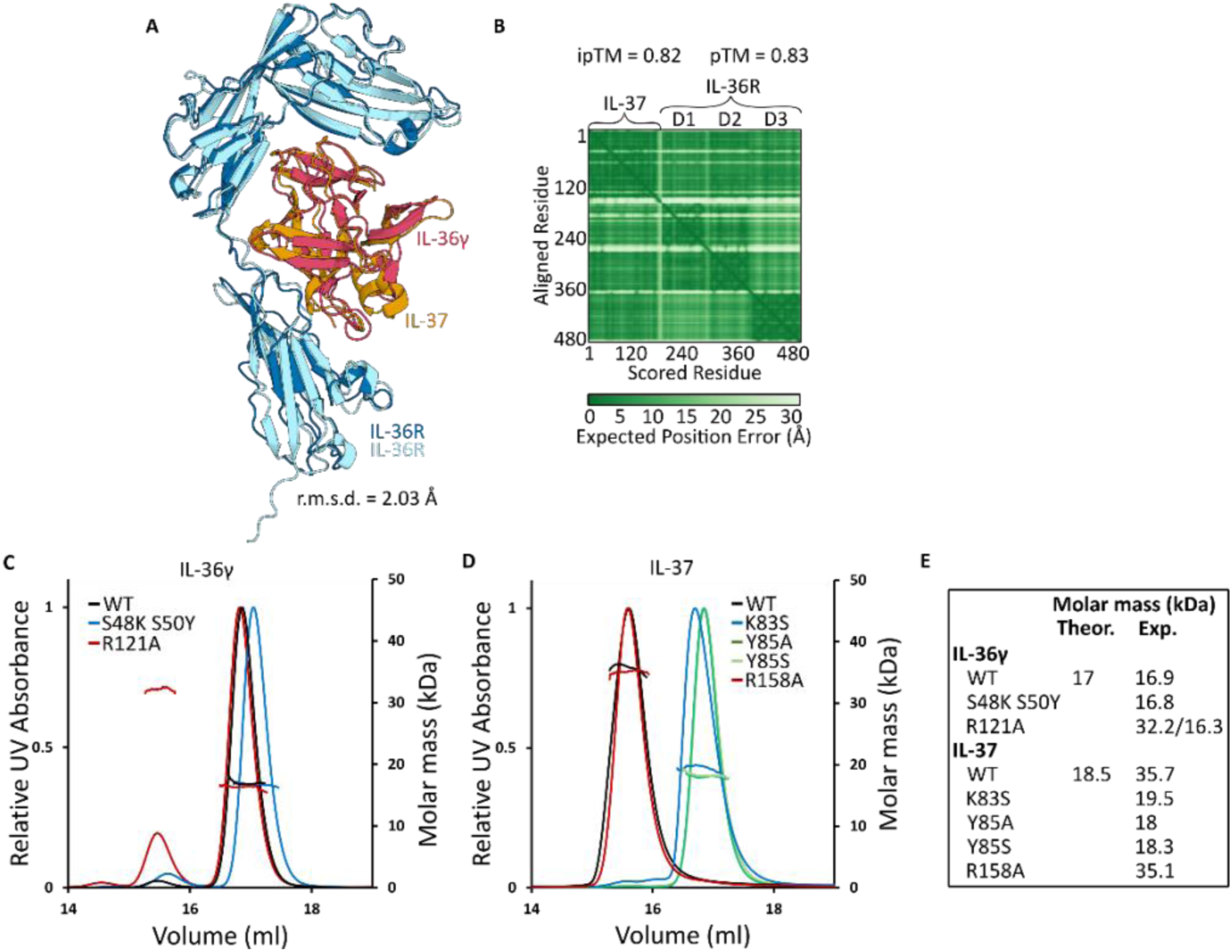
AlphaFold3 predicted IL-37:IL-36R complex used for structure-guided mutagenesis and SEC-MALLS analysis of resulting mutants. (**A**) AlphaFold3 prediction of IL-37 in complex with IL-36R (colored orange/light blue) using the Google DeepMind webserver, aligned with the IL-36γ:IL-36R complex (colored pink/dark blue) from the cryo-EM structure. Alignment was performed with the *align* function in PyMol v2.3.3, and the calculated root mean square deviation (r.m.s.d.) is indicated. (**B**) Predicted aligned error (PAE) plot and (i)pTM scores of the Alphafold3 model. (**C-D**) SEC-MALLS analysis of IL-36γ (C), IL-37 (D), and mutants (C-D) thereof. The SEC-elution profile is plotted as the relative ultraviolet absorbance at 280 nm (left Y-axis) in function of elution volume. The molecular mass as determined by MALLS is plotted on the right y-axis. (**E**) The theoretical (Theor.) and experimental (Exp.) molecular masses are reported. The experimental molecular mass is reported as the average throughout the elution peak. Number of samples analysed: *n* = 1.

### Supplementary tables

**Table S1:**
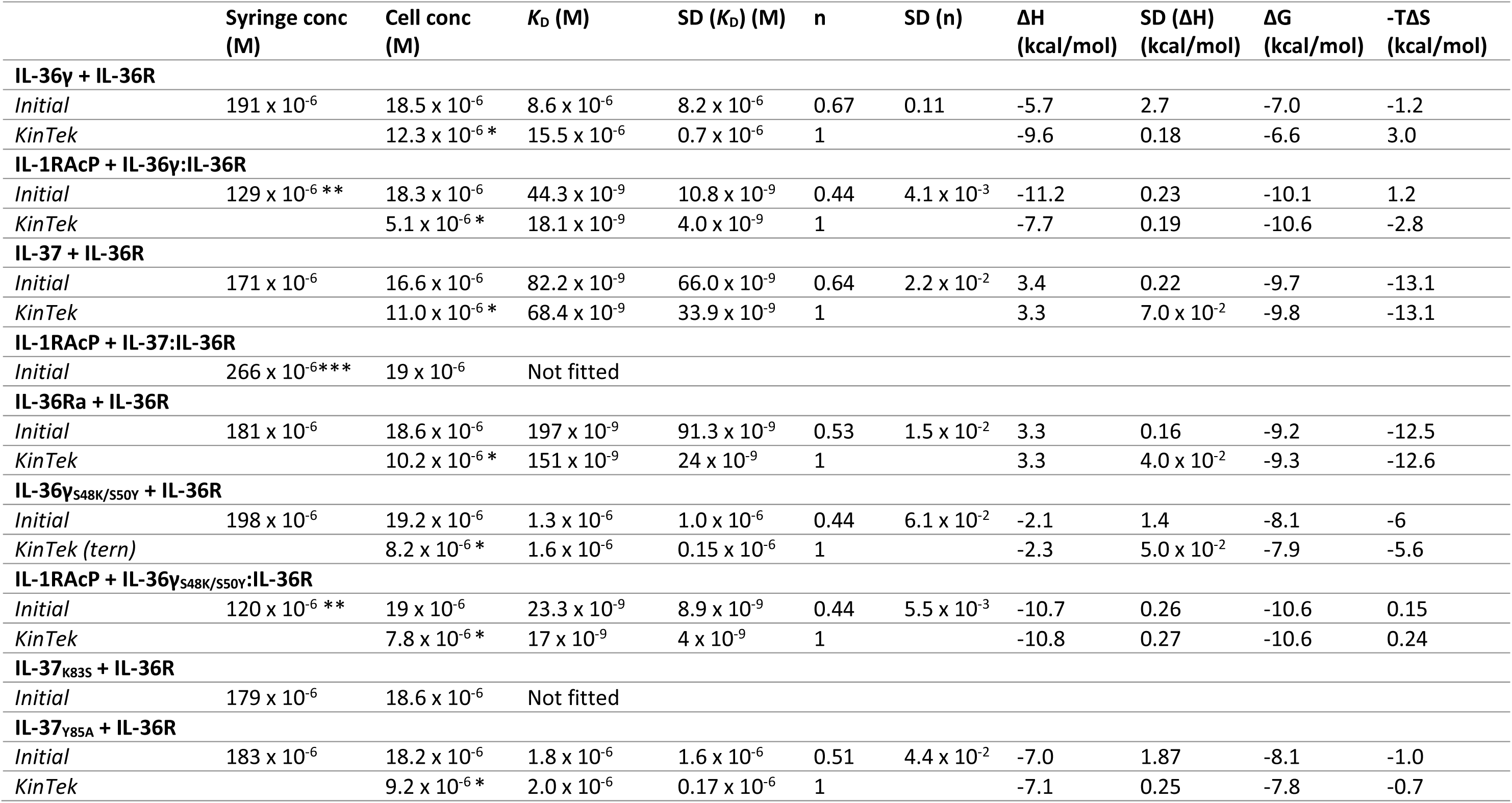
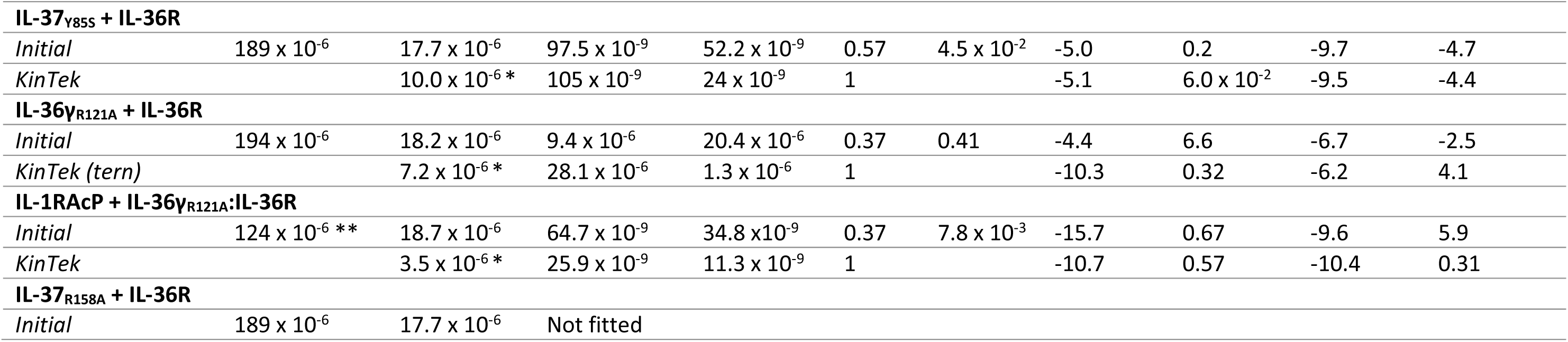
Experimental parameters and derived parameters from ITC experiments. Data was initially analyzed using PEAQ-ITC analysis software (version 1.1.0.1262, Malvern) and fit using a ‘one-set-of-sites’ binding model. To improve the accuracy of the thermodynamic parameters, a global numerical fitting was performed using KinTek Explorer 10 (KinTek Corporation, USA). All values represent an average of at least three replicates. Abbreviations: Conc, concentration, SD, Standard Deviation. *Calculation of active portion of IL-36R. **Experiments performed at approximately 190 and 100 µM, average concentration is reported. *** Experiments performed at approximately 190 and 427 µM, average concentration is reported.

**Table S2:** Cryo-EM data collection, refinement and validation statistics.

|  |  |
| --- | --- |
|  | IL-36γ:IL-36R:IL-1RAP<br>EMDB code: EMD-52665<br>PDB code: 9I7X |
| **Data collection and processing** |  |
| Magnification | 60,000 |
| Voltage (kV) | 300 |
| Electron exposure (e<sup>-</sup> Å<sup>-2</sup>) | 61.8 |
| Defocus range (μm) | 0.8-1.7 |
| Pixel size (Å) | 0.74 |
| Symmetry Imposed | C1 |
| Number of initial particle images | 811,083 |
| Number of final particle images | 201,760 |
| Map resolution (Å) | 3.3 |
| FSC threshold | 0.143 |
| Map resolution range (Å) | 6.5 - 2.8 |
| **Refinement** |  |
| Initial model used | AlphaFold2-Multimer models of individual and complexed components |
| Model resolution (Å) | 3.4 |
| FSC threshold | 0.5 |
| Map sharpening *B* factor (Å<sup>2</sup>) | -70 |
| Model composition |  |
| Nonhydrogen atoms | 6251 |
| Protein residues | 765 |
| Ligands | NAG: 11 |
| *B* factors (Å<sup>2</sup>) (min/max/mean) |  |
| Protein | 12.17/198.11/98.81 |
| Ligand | 83.38/170.50/133.51 |
| R.m.s.d. |  |
| Bond lengths (Å) | 0.003 |
| Bond angles (°) | 0.575 |
| **Validation** |  |
| MolProbity score | 1.88 |
| Clash score | 8.63 |
| Poor rotamers (%) | 0.29 |
| CABLAM outliers (%) | 3.93 |
| CC (mask/volume) (sharpened; unsharpened) | 0.86/0.84; 0.84/0.82 |
| EMRinger (sharpened/unsharpened) | 2.51/2.30 |
| Ramachandran plot |  |
| Favored (%) | 93.74 |
| Allowed (%) | 6.26 |
| Disallowed (%) | 0 |
| Ramachandran Z-score |  |
| Whole | -0.90 |
| Helix | 0.05 |
| Sheet | 0.49 |
| Loop | -1.45 |

**Table S3:**
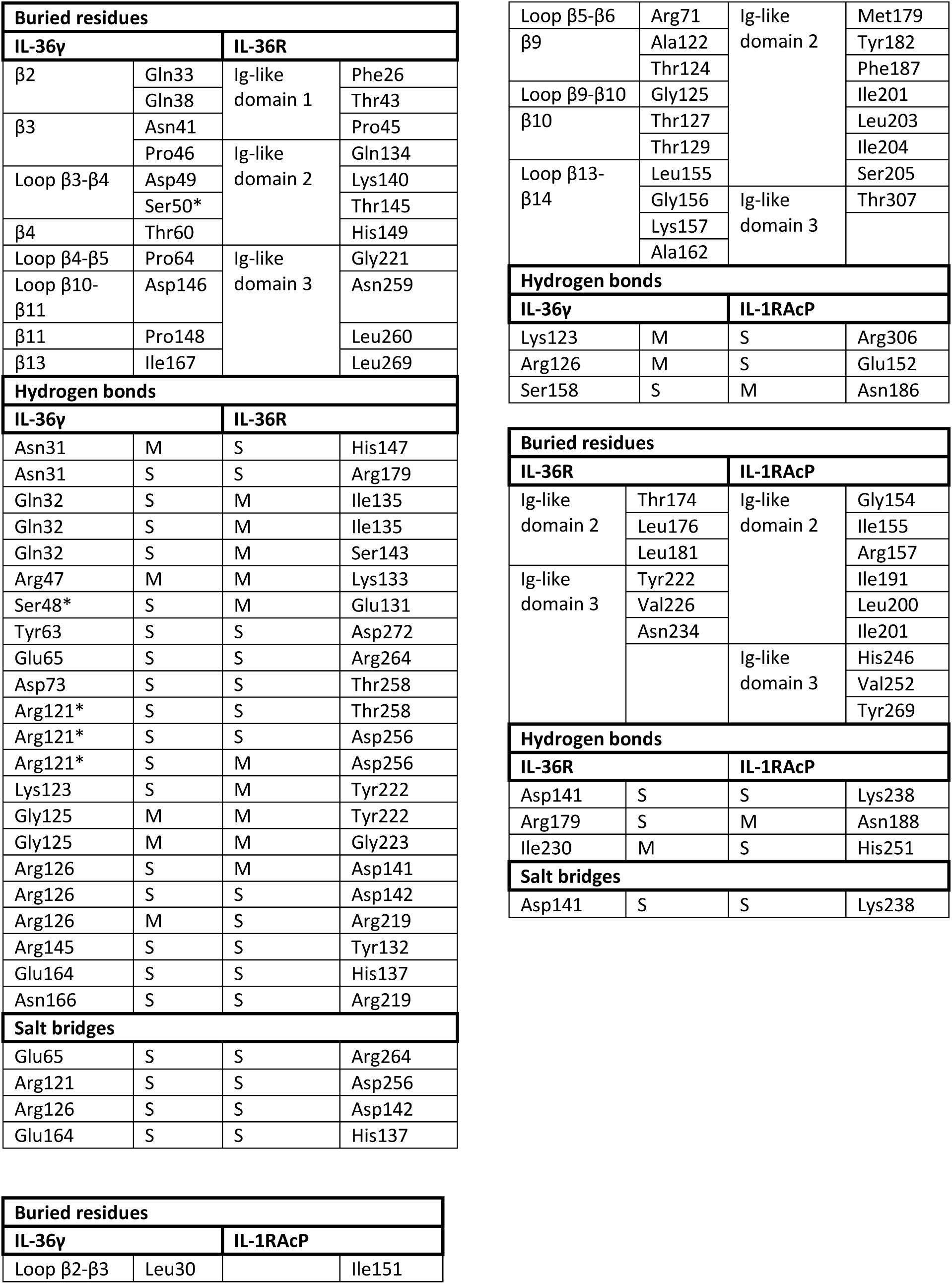
Interaction interface analysis of the IL-36γ:IL-36R:IL-1RAcP ternary complex. Residues that were mutated in this study are marked with an asterisk (*). Involvement of side chain (S) or main chain (M) is indicated for polar interactions. Based on PDBePISA analysis of the cryo-EM structure presented in this study.

**Table S4:** Comparative analysis of the IL-1RAcP interface with human IL-36γ, human IL-1β and mouse IL-33. Based on PDBePISA analysis of the cryo-EM structure presented in this study and published crystal structures of the IL-1β complex (PDB: 4dep) and the mouse IL-33 complex (PDB: 5vi4).

| **Buried residues** |  |  |  |  |  |
| --- | --- | --- | --- | --- | --- |
| **IL-36γ** | **IL-1RAcP** | **IL-1β** | **IL-1RAcP** | **mIL-33** | **mIL-1RAcP** |
| Leu30 | Ile151 | Ile220 | Ile151 | Tyr126 | Phe187 |
| Arg71 | Met179 | Ile222 | Met179 | Lys222 | Leu203 |
| Ala122 | Tyr182 | Lys224 | Tyr182 | Ser223 | Ser205 |
| Thr124 | Phe187 | Thr253 | Phe187 | Met260 | Ser305 |
| Gly125 | Ile201 | Lys254 | Leu203 |  |  |
| Thr127 | Leu203 | Gly255 | Ile204 |  |  |
| Thr129 | Ile204 | Ile259 | Thr307 |  |  |
| Leu155 | Ser205 |  |  |  |  |
| Gly156 | Thr307 |  |  |  |  |
| Lys157 |  |  |  |  |  |
| Ala162 |  |  |  |  |  |
| **Hydrogen bonds** |  |  |  |  |  |
| **IL-36γ** | **IL-1RAcP** | **IL-1β** | **IL-1RAcP** | **mIL-33** | **mIL-1RAcP** |
| Lys123 | Arg306 | Asp170 | Arg306 | Ser224 | Glu152 |
| Arg126 | Glu152 | Asn223 | Glu152 | Asp253 | Lys179 |
| Ser158 | Asn186 | Gln257 | Gln185 | Gln169 | Ser306 |
|  |  | Gln257 | Asn186 | Asp172 | Ser303 |
|  |  | Gln242 | Asn186 | Glu254 | His188 |
|  |  | Gly256 | Asn188 | Tyr126 | His188 |
|  |  | Asp258 | Asn188 | Ser256 | His188 |
|  |  | Ser129 | Asn188 |  |  |
|  |  | Gln130 | Asn188 |  |  |
|  |  | Asp261 | Ser205 |  |  |
| **Salt bridges** |  |  |  |  |  |
| **IL-36γ** | **IL-1RAcP** | **IL-1β** | **IL-1RAcP** | **mIL-33** | **mIL-1RAcP** |
|  |  | Asp170 | Arg306 | Asp253 | Lys179 |

**Table S5:** Comparative analysis of the IL-1RAcP interface with human IL-36R, human IL-1R1 and mouse ST2. Based on PDBePISA analysis of the cryo-EM structure presented in this study and published crystal structures of the IL-1β complex (PDB: 4dep) and the mouse IL-33 complex (PDB: 5vi4).

| **Buried residues** |  |  |  |  |  |
| --- | --- | --- | --- | --- | --- |
| **IL-36R** | **IL-1RAcP** | **IL-1R1** | **IL-1RAcP** | **mST2** | **mIL-1RAcP** |
| Thr174 | Gly154 | Val177 | Ile155 | Phe135 | Gly154 |
| Leu176 | Ile155 | Ile182 | Ile191 | Tyr171 | His188 |
| Leu181 | Arg157 | Met185 | Leu200 | Phe173 | Asn189 |
| Tyr222 | Ile191 | Pro223 | Ile201 | Phe213 | Leu191 |
| Val226 | Leu200 | His318 | Asn239 | Val218 | Phe200 |
| Asn234 | Ile201 |  | Val252 | Thr220 | Ile201 |
|  | His246 |  | Thr307 | Pro223 | Leu241 |
|  | Val252 |  | Thr311 | Lys246 | Tyr246 |
|  | Tyr269 |  |  |  | Tyr269 |
| **Hydrogen bonds** |  |  |  |  |  |
| **IL-36R** | **IL-1RAcP** | **IL-1R1** | **IL-1RAcP** | **mST2** | **mIL-1RAcP** |
| Asp141 | Lys238 | Asp137 | Lys238 | Glu210 | Lys238 |
| Arg179 | Asn188 | Arg180 | Asn188 | Asp175 | Ile155 |
| Ile230 | His251 | Val229 | His251 | Asp284 | Gln244 |
|  |  | Asp137 | Gly154 | Asn221 | Asn249 |
|  |  | Asn185 | Lys238 | Arg169 | Asp186 |
|  |  | Asp321 | His246 | His168 | Phe187 |
|  |  | Arg225 | Tyr269 | His168 | Val190 |
|  |  |  |  | Arg166 | Glu193 |
|  |  |  |  | Arg166 | Ser198 |
|  |  |  |  | Lys133 | Asp239 |
|  |  |  |  | Met285 | Gln244 |
|  |  |  |  | Asn283 | Ile245 |
| **Salt bridges** |  |  |  |  |  |
| **IL-36R** | **IL-1RAcP** | **IL-1R1** | **IL-1RAcP** | **mST2** | **mIL-1RAcP** |
| Asp141 | Lys238 | Asp137 | Lys238 | Glu210 | Lys238 |
|  |  | Asp321 | His246 | Arg169 | Asp186 |
|  |  |  |  | His168 | Asp186 |
|  |  |  |  | Arg166 | Glu193 |
|  |  |  |  | Lys133 | Asp239 |

## References and Notes

1. L. Mercurio, C. M. Failla, L. Capriotti, C. Scarponi, F. Facchiano, M. Morelli, S. Rossi, G. Pagnanelli, C. Albanesi, A. Cavani, S. Madonna, Interleukin (IL)-17/IL-36 axis participates to the crosstalk between endothelial cells and keratinocytes during inflammatory skin responses. PLoS One 15 (2020).

2. S. Madonna, G. Girolomoni, C. A. Dinarello, C. Albanesi, The significance of il-36 hyperactivation and il-36r targeting in psoriasis. Int J Mol Sci 20 (2019).

3. K. L. Sachen, C. N. Arnold Greving, J. E. Towne, Role of IL-36 cytokines in psoriasis and other inflammatory skin conditions. Cytokine 156 (2022).

4. J. E. Towne, K. E. Garka, B. R. Renshaw, G. D. Virca, J. E. Sims, Interleukin (IL)-1F6, IL-1F8, and IL-1F9 Signal Through IL-1Rrp2 and IL-1RAcP to Activate the Pathway Leading to NF-κB and MAPKs. Journal of Biological Chemistry 279, 13677–13688 (2004).

5. R. Debets, J. C. Timans, B. Homey, S. Zurawski, T. R. Sana, S. Lo, J. Wagner, G. Edwards, T. Clifford, S. Menon, J. F. Bazan, R. A. Kastelein, Two Novel IL-1 Family Members, IL-1δ and IL-1ε, Function as an Antagonist and Agonist of NF-B Activation Through the Orphan IL-1 Receptor-Related Protein 2 1. The Journal of Immunology 167, 1440–1446 (2001).

6. F. L. Van De Veerdonk, A. K. Stoeckman, G. Wu, A. N. Boeckermann, T. Azam, M. G. Netea, L. A. B. Joosten, J. W. M. Van Der Meer, R. Hao, V. Kalabokis, C. A. Dinarello, IL-38 binds to the IL-36 receptor and has biological effects on immune cells similar to IL-36 receptor antagonist. Proceedings of the National Academy of Sciences 109, 3001–3005 (2012).

7. A. M. Foster, J. Baliwag, C. S. Chen, A. M. Guzman, S. W. Stoll, J. E. Gudjonsson, N. L. Ward, A. Johnston, IL-36 Promotes Myeloid Cell Infiltration, Activation, and Inflammatory Activity in Skin. The Journal of Immunology 192, 6053–6061 (2014).

8. D. Dietrich, P. Martin, V. Flacher, Y. Sun, D. Jarrossay, N. Brembilla, C. Mueller, H. A. Arnett, G. Palmer, J. Towne, C. Gabay, Interleukin-36 potently stimulates human M2 macrophages, Langerhans cells and keratinocytes to produce pro-inflammatory cytokines. Cytokine 84, 88–98 (2016).

9. A. Onoufriadis, M. A. Simpson, A. E. Pink, P. Di Meglio, C. H. Smith, V. Pullabhatla, J. Knight, S. L. Spain, F. O. Nestle, A. D. Burden, F. Capon, R. C. Trembath, J. N. Barker, Mutations in IL36RN/IL1F5 Are Associated with the Severe Episodic Inflammatory Skin Disease Known as Generalized Pustular Psoriasis. Am J Hum Genet 89, 432 (2011).

10. N. Setta-Kaffetzi, A. A. Navarini, V. M. Patel, V. Pullabhatla, A. E. Pink, S. E. Choon, M. A. Allen, A. D. Burden, C. E. M. Griffiths, M. M. B. Seyger, B. Kirby, R. C. Trembath, M. A. Simpson, C. H. Smith, F. Capon, J. N. Barker, Rare Pathogenic Variants in IL36RN Underlie a Spectrum of Psoriasis-Associated Pustular Phenotypes. Journal of Investigative Dermatology 133, 1366–1369 (2013).

11. S. J. Martin, V. Frezza, P. Davidovich, Z. Najda, D. M. Clancy, IL-1 family cytokines serve as “activity recognition receptors” for aberrant protease activity indicative of danger. Cytokine 157 (2022).

12. I. S. Afonina, C. Müller, S. J. Martin, R. Beyaert, Proteolytic Processing of Interleukin-1 Family Cytokines: Variations on a Common Theme. Immunity 42, 991–1004 (2015).

13. J. E. Towne, B. R. Renshaw, J. Douangpanya, B. P. Lipsky, M. Shen, C. A. Gabel, J. E. Sims, Interleukin-36 (IL-36) Ligands Require Processing for Full Agonist (IL-36, IL-36, and IL-36) or Antagonist (IL-36Ra) Activity □ S. doi: 10.1074/jbc.M111.267922 (2011).

14. C. M. Henry, G. P. Sullivan, D. M. Clancy, I. S. Afonina, D. Kulms, S. J. Martin, Neutrophil-Derived Proteases Escalate Inflammation through Activation of IL-36 Family Cytokines. Cell Rep 14, 708–722 (2016).

15. S. J. Busfield, C. A. Comrack, G. Yu, T. W. Chickering, J. S. Smutko, H. Zhou, K. R. Leiby, L. M. Holmgren, D. P. Gearing, Y. Pan, Identification and Gene Organization of Three Novel Members of the IL-1 Family on Human Chromosome 2. Genomics 66, 213–216 (2000).

16. D. E. Smith, B. R. Renshaw, R. R. Ketchem, M. Kubin, K. E. Garka, J. E. Sims, Four new members expand the interleukin-1 superfamily. J Biol Chem 275, 1169–1175 (2000).

17. S. Kumar, P. C. McDonnell, R. Lehr, L. Tierney, M. N. Tzimas, D. E. Griswold, E. A. Capper, R. Tal-Singer, G. I. Wells, M. L. Doyle, P. R. Young, Identification and initial characterization of four novel members of the interleukin-1 family. J Biol Chem 275, 10308–10314 (2000).

18. G. Pan, P. Risser, W. Mao, D. T. Baldwin, A. W. Zhong, E. Filvaroff, D. Yansura, L. Lewis, C. Eigenbrot, W. J. Henzel, R. Vandlen, IL-1H, AN INTERLEUKIN 1-RELATED PROTEIN THAT BINDS IL-18 RECEPTOR/IL-1Rrp. Cytokine 13, 1–7 (2001).

19. C. A. Nold-Petry, C. Y. Lo, I. Rudloff, K. D. Elgass, S. Li, M. P. Gantier, A. S. Lotz-Havla, S. W. Gersting, S. X. Cho, J. C. Lao, A. M. Ellisdon, B. Rotter, T. Azam, N. E. Mangan, F. J. Rossello, J. C. Whisstock, P. Bufler, C. Garlanda, A. Mantovani, C. A. Dinarello, M. F. Nold, IL-37 requires the receptors IL-18Rα and IL-1R8 (SIGIRR) to carry out its multifaceted anti-inflammatory program upon innate signal transduction. Nature Immunology 2015 16:4 16, 354–365 (2015).

20. G. P. Sullivan, P. Davidovich, N. Muñoz-Wolf, R. W. Ward, Y. E. Hernandez Santana, D. M. Clancy, A. Gorman, Z. Najda, B. Turk, P. T. Walsh, E. C. Lavelle, S. J. Martin, Myeloid cell–derived proteases produce a proinflammatory form of IL-37 that signals via IL-36 receptor engagement. Sci Immunol 7 (2022).

21. S. Kumar, C. R. Hanning, M. R. Brigham-Burke, D. J. Rieman, R. Lehr, S. Khandekar, R. B. Kirkpatrick, G. F. Scott, J. C. Lee, F. J. Lynch, W. Gao, A. Gambotto, M. T. Lotze, INTERLEUKIN-1F7B (IL-1H4/IL-1F7) IS PROCESSED BY CASPASE-1 AND MATURE IL-1F7B BINDS TO THE IL-18 RECEPTOR BUT DOES NOT INDUCE IFN-gamma PRODUCTION. Cytokine 18, 61–71 (2002).

22. S. Sharma, N. Kulk, M. F. Nold, R. Gräf, S.-H. Kim, D. Reinhardt, C. A. Dinarello, P. Bufler, The IL-1 Family Member 7b Translocates to the Nucleus and Down-Regulates Proinflammatory Cytokines 1. The Journal of Immunology 180, 5477–5482 (2008).

23. A. Morita, B. Strober, D. Burden, S. E. Choon, M. J. Anadkat, S. Marrakchi, T.-F. Tsai, K. B. Gordon, D. Thaçi, M. Zheng, N. Hu, T. Haeufel, C. Thoma, M. G. Lebwohl, Efficacy and safety of subcutaneous spesolimab for the prevention of generalised pustular psoriasis flares (Effisayil 2): an international, multicentre, randomised, placebo-controlled trial. www.thelancet.com 402, 2023 (2023).

24. Boehringer Ingelheim, SPEVIGO® approved for expanded indications in China and the US (2024). https://www.boehringer-ingelheim.com/us/human-health/skin-and-inflammatory-diseases/gpp/spevigo-approved-expanded-indications-china-and-us.

25. R. Ganesan, E. L. Raymond, D. Mennerich, J. R. Woska, G. Caviness, C. Grimaldi, J. Ahlberg, R. Perez, S. Roberts, D. Yang, K. Jerath, K. Truncali, L. Frego, E. Sepulveda, P. Gupta, S. E. Brown, M. D. Howell, K. A. Canada, R. Kroe-Barrett, J. S. Fine, S. Singh, M. L. Mbow, Generation and functional characterization of anti-human and anti-mouse IL-36R antagonist monoclonal antibodies. MAbs 9, 1143–1154 (2017).

26. E. T. Larson, D. L. Brennan, E. R. Hickey, R. Ganesan, R. Kroe-Barrett, N. A. Farrow, X-ray crystal structure localizes the mechanism of inhibition of an IL-36R antagonist monoclonal antibody to interaction with Ig1 and Ig2 extra cellular domains. Protein Science 29, 1679–1686 (2020).

27. J. Hou, S. A. Townson, J. T. Kovalchin, A. Masci, O. Kiner, Y. Shu, B. M. King, E. Schirmer, K. Golden, C. Thomas, K. C. Garcia, G. Zarbis-Papastoitsis, E. S. Furfine, T. M. Barnes, Design of a superior cytokine antagonist for topical ophthalmic use. CELL BIOLOGY 110, 3913–3918 (2013).

28. S. Detry, J. Andries, Y. Bloch, C. Gabay, D. M. Clancy, S. N. Savvides, Structural basis of human IL-18 sequestration by the decoy receptor IL-18 binding protein in inflammation and tumor immunity. Journal of Biological Chemistry 298, 101908 (2022).

29. X. Liu, M. Hammel, Y. He, J. A. Tainer, U. S. Jeng, L. Zhang, S. Wang, X. Wang, Structural insights into the interaction of IL-33 with its receptors. Proc Natl Acad Sci U S A 110, 14918–14923 (2013).

30. Zhou, V. Todorovic, S. Kakavas, B. Sielaff, L. Medina, L. Wang, R. Sadhukhan, H. Stockmann, P. L. Richardson, E. DiGiammarino, C. Sun, V. Scott, Quantitative ligand and receptor binding studies reveal the mechanism of interleukin-36 (IL-36) pathway activation. Journal of Biological Chemistry 293, 403–411 (2018).

31. K. A. Johnson, Chapter 23 Fitting Enzyme Kinetic Data with KinTek Global Kinetic Explorer. Methods Enzymol 467, 601–626 (2009).

32. K. A. Johnson, Z. B. Simpson, T. Blom, Global Kinetic Explorer: A new computer program for dynamic simulation and fitting of kinetic data. Anal Biochem 387, 20–29 (2009).

33. K. A. Johnson, Z. B. Simpson, T. Blom, FitSpace Explorer: An algorithm to evaluate multidimensional parameter space in fitting kinetic data. Anal Biochem 387, 30–41 (2009).

34. D. M. Clancy, C. M. Henry, G. P. Sullivan, S. J. Martin, Neutrophil extracellular traps can serve as platforms for processing and activation of IL-1 family cytokines. FEBS J 284, 1712–1725 (2017).

35. S. Günther, E. J. Sundberg, Molecular Determinants of Agonist and Antagonist Signaling through the IL-36 Receptor. The Journal of Immunology 193, 921–930 (2014).

36. C. Thomas, J. F. Bazan, K. C. Garcia, Structure of the activating IL-1 receptor signaling complex. Nat Struct Mol Biol 19, 455 (2012).

37. N. Tsutsumi, T. Kimura, K. Arita, M. Ariyoshi, H. Ohnishi, T. Yamamoto, X. Zuo, K. Maenaka, E. Y. Park, N. Kondo, M. Shirakawa, H. Tochio, Z. Kato, The structural basis for receptor recognition of human interleukin-18. Nat Commun 5 (2014).

38. D. Wang, S. Zhang, L. Li, X. Liu, K. Mei, X. Wang, Structural insights into the assembly and activation of IL-1β with its receptors. Nature Immunology 2010 11:10 11, 905–911 (2010).

39. S. Günther, D. Deredge, A. L. Bowers, A. Luchini, D. A. Bonsor, R. Beadenkopf, L. Liotta, P. L. Wintrode, E. J. Sundberg, IL-1 Family Cytokines Use Distinct Molecular Mechanisms to Signal through Their Shared Co-receptor. Immunity 47, 510–523.e4 (2017).

40. E. Krissinel, K. Henrick, Inference of Macromolecular Assemblies from Crystalline State. J. Mol. Biol. 372, 774–797 (2007).

41. S. Marrakchi, P. Guigue, B. R. Renshaw, A. Puel, X.-Y. Pei, S. Fraitag, J. Zribi, E. Bal, C. Cluzeau, M. Chrabieh, J. E. Towne, J. Douangpanya, C. Pons, S. Mansour, V. Serre, H. Makni, N. Mahfoudh, F. Fakhfakh, C. Bodemer, J. Feingold, S. Hadj-Rabia, M. Favre, E. Genin, M. Sahbatou, A. Munnich, J.-L. Casanova, J. E. Sims, H. Turki, H. Bachelez, A. Smahi, Interleukin-36–Receptor Antagonist Deficiency and Generalized Pustular Psoriasis. New England Journal of Medicine 365, 620–628 (2011).

42. A. M. Ellisdon, C. A. Nold-Petry, L. D’Andrea, S. X. Cho, J. C. Lao, I. Rudloff, D. Ngo, C. Y. Lo, T. P. Soares da Costa, M. A. Perugini, P. J. Conroy, J. C. Whisstock, M. F. Nold, Homodimerization attenuates the anti-inflammatory activity of interleukin-37. Sci Immunol 2 (2017).

43. S. Il Yoon, B. C. Jones, N. J. Logsdon, M. R. Walter, Same Structure, Different Function: Crystal Structure of the Epstein-Barr Virus IL-10Bound to the Soluble IL-10R1 Chain. Structure 13, 551–564 (2005).

44. E. Z. Eisenmesser, A. Gottschlich, J. S. Redzic, N. Paukovich, J. C. Nix, T. Azam, L. Zhang, R. Zhao, J. S. Kieft, E. The, X. Meng, C. A. Dinarello, Interleukin-37 monomer is the active form for reducing innate immunity. Proceedings of the National Academy of Sciences 116, 5514–5522 (2019).

45. M. Tauber, E. Bal, X. Y. Pei, M. Madrange, A. Khelil, H. Sahel, A. Zenati, M. Makrelouf, K. Boubridaa, A. Chiali, N. Smahi, F. Otsmane, B. Bouajar, S. Marrakchi, H. Turki, E. Bourrat, M. Viguier, Y. Hamel, H. Bachelez, A. Smahi, IL36RN Mutations Affect Protein Expression and Function: A Basis for Genotype-Phenotype Correlation in Pustular Diseases. Journal of Investigative Dermatology 136, 1811–1819 (2016).

46. G. P. A. Vigers, L. J. Anderson, P. Caffes, B. J. Brandhuber, Crystal structure of the type-I interleukin-1 receptor complexed with interleukin-1β. Nature 1997 386:6621 386, 190–194 (1997).

47. Z. C. Yuan, W. D. Xu, X. Y. Liu, X. Y. Liu, A. F. Huang, L. C. Su, Biology of il-36 signaling and its role in systemic inflammatory diseases. Front Immunol 10, 467377 (2019).

48. D. Queen, C. Ediriweera, L. Liu, Function and Regulation of IL-36 Signaling in Inflammatory Diseases and Cancer Development. Front Cell Dev Biol 7 (2019).

49. M. J. Gooderham, A. S. Van Voorhees, M. G. Lebwohl, An update on generalized pustular psoriasis. Expert Rev Clin Immunol 15, 907–919 (2019).

50. J. K. Fields, E. J. Gyllenbäck, M. Bogacz, J. Obi, G. S. Birkedal, K. Sjöström, K. Maravillas, C. Grönberg, S. Rattik, K. Kihn, M. Flowers, A. K. Smith, N. Hansen, T. Fioretos, C. Huyhn, D. Liberg, D. Deredge, E. J. Sundberg, Antibodies targeting the shared cytokine receptor IL-1 receptor accessory protein invoke distinct mechanisms to block all cytokine signaling. Cell Rep 0, 114099 (2024).

51. Y. Dong, J. P. Bonin, P. Devant, Z. Liang, A. I. M. Sever, J. Mintseris, J. M. Aramini, G. Du, S. P. Gygi, J. C. Kagan, L. E. Kay, H. Wu, Structural transitions enable interleukin-18 maturation and signaling. Immunity 57, 1533–1548.e10 (2024).

52. P. Devant, Y. Dong, J. Mintseris, W. Ma, S. P. Gygi, H. Wu, J. C. Kagan, Structural insights into cytokine cleavage by inflammatory caspase-4. Nature 624, 451–459 (2023).

53. D. M. Clancy, J. Andries, S. N. Savvides, The pros and confs of IL-18 activation. Immunity 57, 1445–1448 (2024).

54. A. R. Aricescu, W. Lu, E. Y. Jones, A time- and cost-efficient system for high-level protein production in mammalian cells. Acta Crystallogr D Biol Crystallogr 62, 1243–1250 (2006).

55. D. N. Mastronarde, Automated electron microscope tomography using robust prediction of specimen movements. J Struct Biol 152, 36–51 (2005).

56. A. Punjani, J. L. Rubinstein, D. J. Fleet, M. A. Brubaker, cryosPArc: algorithms for rapid unsupervised cryo-em structure determination. Nat Methods 14, 290–297 (2017).

57. T. Wagner, F. Merino, M. Stabrin, T. Moriya, C. Antoni, A. Apelbaum, P. Hagel, O. Sitsel, T. Raisch, D. Prumbaum, D. Quentin, D. Roderer, S. Tacke, B. Siebolds, E. Schubert, T. R. Shaikh, P. Lill, C. Gatsogiannis, S. Raunser, SPHIRE-crYOLO is a fast and accurate fully automated particle picker for cryo-EM. Nature Communications Biology 2 (2019).

58. T. Bepler, A. Morin, M. Rapp, J. Brasch, L. Shapiro, A. J. Noble, B. Berger, Positive-unlabeled convolutional neural networks for particle picking in cryo-electron micrographs. Nat Methods 16, 1153–1160 (2019).

59. P. B. Rosenthal, R. Henderson, Optimal Determination of Particle Orientation, Absolute Hand, and Contrast Loss in Single-particle Electron Cryomicroscopy. J. Mol. Biol. 333, 721–745 (2003).

60. J. Jumper, R. Evans, A. Pritzel, T. Green, M. Figurnov, O. Ronneberger, K. Tunyasuvunakool, R. Bates, A. Žídek, A. Potapenko, A. Bridgland, C. Meyer, S. A. A. Kohl, A. J. Ballard, A. Cowie, B. Romera-Paredes, S. Nikolov, R. Jain, J. Adler, T. Back, S. Petersen, D. Reiman, E. Clancy, M. Zielinski, M. Steinegger, M. Pacholska, T. Berghammer, S. Bodenstein, D. Silver, O. Vinyals, A. W. Senior, K. Kavukcuoglu, P. Kohli, D. Hassabis, D. □ Hassabis, Highly accurate protein structure prediction with AlphaFold. Nature 596, 583 (2021).

61. E. F. Pettersen, T. D. Goddard, C. C. Huang, E. C. Meng, G. S. Couch, T. I. Croll, J. H. Morris, T. E. Ferrin, C. E. Thomas Ferrin, UCSF ChimeraX: Structure visualization for researchers, educators, and developers. Protein Science 30, 70–82 (2020).

62. R. T. Kidmose, J. Juhl, P. Nissen, T. Boesen, J. L. Karlsen, B. P. Pedersen, Namdinator-automatic molecular dynamics flexible fitting of structural models into cryo-EM and crystallography experimental maps. IUCrJ 6, 526–531 (2019).

63. P. Emsley, K. Cowtan, Biological Crystallography Coot: model-building tools for molecular graphics. Acta Crystallographica Section D D60, 2126–2132 (2004).

64. D. Liebschner, P. V Afonine, M. L. Baker, G. Bunkó, V. B. Chen, T. I. Croll, B. Hintze, L.-W. Hung, S. Jain, } Airlie, J. Mccoy, N. W. Moriarty, R. D. Oeffner, B. K. Poon, M. G. Prisant, R. J. Read, J. S. Richardson, D. C. Richardson, M. D. Sammito, O. V Sobolev, D. H. Stockwell, T. C. Terwilliger, A. G. Urzhumtsev, L. L. Videau, C. J. Williams, P. D. Adams, Macromolecular structure determination using X-rays, neutrons and electrons: recent developments in Phenix. Acta Cryst 75, 861–877 (2019).

65. C. J. Williams, J. J. Headd, N. W. Moriarty, M. G. Prisant, L. L. Videau, L. N. Deis, V. Verma, D. A. Keedy, B. J. Hintze, V. B. Chen, S. Jain, S. M. Lewis, W. B. Arendall, J. Snoeyink, P. D. Adams, S. C. Lovell, J. S. Richardson, D. C. Richardson, MolProbity: More and better reference data for improved all-atom structure validation. Protein Sci 27, 293–315 (2018).

66. J. Abramson, J. Adler, J. Dunger, R. Evans, T. Green, A. Pritzel, O. Ronneberger, L. Willmore, A. J. Ballard, J. Bambrick, S. W. Bodenstein, D. A. Evans, C.-C. Hung, M. O’Neill, D. Reiman, K. Tunyasuvunakool, Z. Wu, A. Žemgulytė, E. Arvaniti, C. Beattie, O. Bertolli, A. Bridgland, A. Cherepanov, M. Congreve, A. I. Cowen-Rivers, A. Cowie, M. Figurnov, F. B. Fuchs, H. Gladman, R. Jain, Y. A. Khan, C. M. R. Low, K. Perlin, A. Potapenko, P. Savy, S. Singh, A. Stecula, A. Thillaisundaram, C. Tong, S. Yakneen, E. D. Zhong, M. Zielinski, A. Žídek, V. Bapst, P. Kohli, M. Jaderberg, D. Hassabis, J. M. Jumper, Accurate structure prediction of biomolecular interactions with AlphaFold 3. Nature 630, 493–500 (2024).

67. F. Sievers, A. Wilm, D. Dineen, T. J. Gibson, K. Karplus, W. Li, R. Lopez, H. McWilliam, M. Remmert, J. Söding, J. D. Thompson, D. G. Higgins, Fast, scalable generation of high-quality protein multiple sequence alignments using Clustal Omega. Mol Syst Biol 7, 539 (2011).

68. X. Robert, P. Gouet, Deciphering key features in protein structures with the new ENDscript server. Nucleic Acids Res 42, W320–W324 (2014).

69. E. Gasteiger, C. Hoogland, A. Gattiker, S. Duvaud, M. R. Wilkins, R. D. Appel, A. Bairoch, Protein Identification and Analysis Tools on the ExPASy Server. The Proteomics Protocols Handbook 52, 571–607 (2005).

70. S. Chen, G. McMullan, A. R. Faruqi, G. N. Murshudov, J. M. Short, S. H. W. Scheres, R. Henderson, High-resolution noise substitution to measure overfitting and validate resolution in 3D structure determination by single particle electron cryomicroscopy. Ultramicroscopy 135, 24–35 (2013).

